# Evolutionary progression of collective mutations in Omicron sub-lineages towards efficient RBD-hACE2: allosteric communications between and within viral and human proteins

**DOI:** 10.1101/2022.08.06.503019

**Authors:** Victor Barozi, Adrienne L. Edkins, Özlem Tastan Bishop

**Author notes:** Corresponding author: Özlem Tastan Bishop.

## Abstract

The interaction between the Spike (S) protein of SARS-CoV-2 and the human angiotensin converting enzyme 2 (hACE2) is essential for infection, and is a target for neutralizing antibodies. Consequently, selection of mutations in the S protein is expected to be driven by the impact on the interaction with hACE2 and antibody escape. Here, for the first time, we systematically characterized the collective effects of mutations in each of the Omicron sub-lineages (BA.1, BA.2, BA.3 and BA.4) on both the viral S protein receptor binding domain (RBD) and the hACE2 protein using post molecular dynamics studies and dynamic residue network (DRN) analysis. Our analysis suggested that Omicron sub-lineage mutations result in altered physicochemical properties that change conformational flexibility compared to the reference structure, and may contribute to antibody escape. We also observed changes in the hACE2 substrate binding groove in some sub-lineages. Notably, we identified unique allosteric communication paths in the reference protein complex formed by the DRN metrics *betweenness centrality* and *eigencentrality* hubs, originating from the RBD core traversing the receptor binding motif of the S protein and the N-terminal domain of the hACE2 to the active site. We showed allosteric changes in residue network paths in both the RBD and hACE2 proteins due to Omicron sub-lineage mutations. Taken together, these data suggest progressive evolution of the Omicron S protein RBD in sub-lineages towards a more efficient interaction with the hACE2 receptor which may account for the increased transmissibility of Omicron variants.

## Introduction

Coronavirus disease-2019, COVID-19, is caused by the severe acute respiratory syndrome coronavirus-2, SARS-CoV-2 [1–3]. It belongs to the *β* genus of the *Coronaviridae* family [4–6]. The other viruses in the *Coronaviridae* family were responsible for the severe acute respiratory syndrome (SARS) in 2002 and the Middle East respiratory syndrome (MERS) in 2012 [7, 8]. The unprecedented “success” in transmission and infectivity of SARS-CoV-2 over the SARS and MERS coronaviruses is a result of evolutionary adaptability through genomic mutations in key regions of the SARS-CoV-2 genome [9–13]. This is evident through the emergence of an extensive number of SARS-CoV-2 variants originating from different geographical locations. The PANGO lineage classification [14] shows the a diverse distribution of SARS-CoV-2 variants across the African continent with the most prevalent being the Omicron variant. The World Health Organization (WHO) classifies these variants according to severity i.e., variants of concern (VOC) with mutations leading to increased transmissibility, virulence and reduced effectiveness of social and drug therapeutics. VOC include the Alpha (B.1.1.7), Beta (B.1.351), Gamma (P.1), Delta (B.1.617.2) and, most prevalently, the Omicron variant (B.1.1.529) [15–18]. These variants contain several single nucleotide polymorphisms (SNPs) and deletions in key SARS-CoV-2 viral proteins facilitating higher receptor affinity, vaccine and neutralizing antibody escape [19–23].

The spike (S) protein is required for SARS-CoV-2 infection through recognition and binding of the host human angiotensin converting enzyme 2 (hACE2) receptor [24–28]. The S protein is a homo-trimer of the S1 and S2 subunits responsible for receptor binding and membrane fusion, respectively [29]. The S1 subunit is composed of the N-terminal domain (NTD; 14-305) and the receptor binding domain (RBD; 319-541) which directly interacts with the hACE2 facilitating SARS-CoV-2 host cell binding (**Figure 1**). All the S protein interactions except Lys417, which forms salt-bridge interaction with Asp30 of hACE2, occur via the receptor binding motif (RBM) of the RBD (**Figure 1**). This motif encompasses residues 438 and 506 in the S protein [6]. The S2 subunit contains the fusion peptide, heptad repeat sequence 1 (HR1), HR2, transmembrane domain (TM) and cytoplasm domain (CM) of the S protein [6, 29, 30].

**Figure 1:**
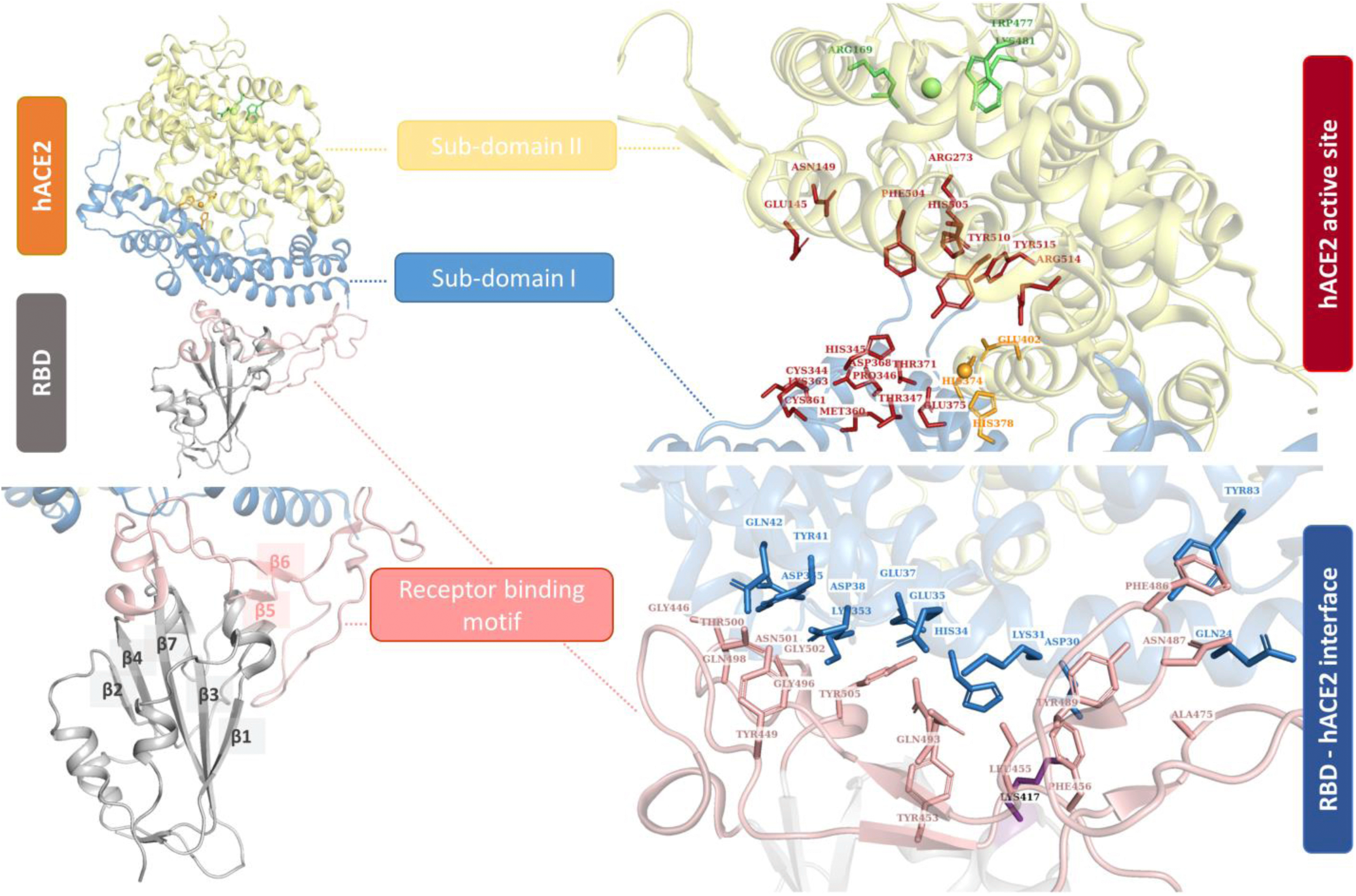
Cartoon representation of the RBD-hACE2 complex (PDB ID: 6M0J) [6] showing the receptor binding motif (RBM) of the RBD in boron and the hACE2 sub-domains I and II in blue and yellow, respectively. hACE2 active site residues are shown as red sticks whereas the zinc ion (orange sphere) and chlorine ion (green sphere) coordinating residues are shown as orange and green sticks, respectively. RBD and hACE2 interface residues involved in complex interaction are shown as boron and sky-blue sticks, respectively. Interface and active site residue data are taken from [6, 35].

ACE2 is a mono-carboxypeptidase [31], that inactivates angiotensin peptides I and II to inhibit the renin-angiotensin system (RAS) which regulates blood pressure [32, 33]. Consequently, ACE2 activity is linked to cardiovascular function and hypertension [34]. The extracellular ACE2 domain, which encompasses the S protein binding site, is predominantly alpha helical, and made up of two domains, a larger N-terminal domain (spanning residues 9 – 611) followed by a smaller C-terminal collectrin homology domain (spanning residues 612-740). The enzyme activity resides in the N-terminal domain, which is divided into catalytic sub-domains I (encompassing residues 19–102, 290–397, and 417–430) and II (encompassing residues 103–289, 398–416, and 431–615) [35] (**Figure 1**). The domain has a typical protease structure composed of a deep cleft-like active site formed between subdomains I and II. The zinc ion required for activity is bound within the cleft towards subdomain I, coordinated by residues His374, His378 and Glu402 and water (forming the HEXXH + E motif). The single chloride ion involved in anion regulation [32, 36] is coordinated by residues Arg169, Trp477 and Lys481 from sub-domain II, distal to both the active site zinc and the ligand binding site. Ligand binding by ACE2 results in encapsulation of the ligand within the cleft via a large ∼16° hinge-like movement of subdomain I relative to sub-domain II (which remains largely stationary). Sub-domains I and II contribute residues equally to ligand binding at the active site (**Figure 1**) [35]. Of the active site residues, Arg273 is required for substrate binding, while His345 and His505 are involved in catalysis, with His345 acting as hydrogen bond donor/acceptor during tetrahedral peptide intermediate formation [31]. The core of the SARS-CoV-2 RBD (residues 333 – 526) consists of a 5 stranded twisted beta sheet (β1-4 and 7), connected by short loops and alpha helices (**Figure 1**). The RBM (residues 4438– 506) lies between β4 and β7 forming a concave surface, which accommodates the N-terminal peptidase subdomain I of ACE2. The N-terminal helix of ACE2 (residues 22 – 57) is responsible for the majority of RBM interactions with additional contacts afforded by a small unstructured sequence from residues 351 – 357 [6] (**Figure 1**).

The S protein, particularly the RBD, is a prime target for viral inhibitors and neutralizing antibodies given its role in SARS-CoV-2 infectivity [37–45]. Structural alterations in the SARS-CoV-2 S protein increase the affinity for the hACE2 receptor up to tenfold compared to the corresponding S protein from SARS-CoV [46, 47]. SARS-CoV-2 has acquired multiple mutations in the Omicron NTD and RBD possibly due to suboptimal neutralization from natural or acquired immunity. The initial Omicron variant, B.1.1.529 harbours roughly 30 SNPs, six residue deletions and three residue insertions in the S protein with 15 SNPs in the RBD alone [48]. Mutations in the RBD and NTD of the S1 subunit contribute to escape of neutralizing antibody therapy through impaired antibody binding [49–55]. Specific Omicron RBD mutations linked to neutralizing antibody escape include: G339D [54], S375F [54], K417N [54, 56], N440K [54, 57, 58], G446S [54, 58], L452R [20], S477N [20], T478K [20], F486V [20] and Q493R [54, 57, 58]. Due to multiple RBD mutations, Omicron has a weaker binding affinity for hACE2 compared to the Alpha and Delta variants [59–62] which suggests an evolutionary bargain between binding affinity and neutralizing antibody offset in the Omicron variant.

Progressively, the Omicron variant has acquired new mutations resulting in evolutionary Omicron sub-lineages; BA.1, BA.2, BA.3, BA.4 and BA.5 with unique mutations per sub-lineage (**Figure 2**). For instance the BA.2 sub-lineage harbours the unique S371F, T376A, D405N and R408S mutations in the RBD which are not present in BA.1[63, 64], whereas the RBD mutation G446S is unique to BA.3 compared to BA.2 [65]. These differences in mutations between the Omicron sub-lineages, have been linked to differences in infectivity and antibody neutralizing activity of the sub-lineages. The BA.1 sub-lineage was considered more infectious than BA.2, while BA.3 had the lowest number of cases of the three sub-lineages [66, 67].

**Figure 2:**
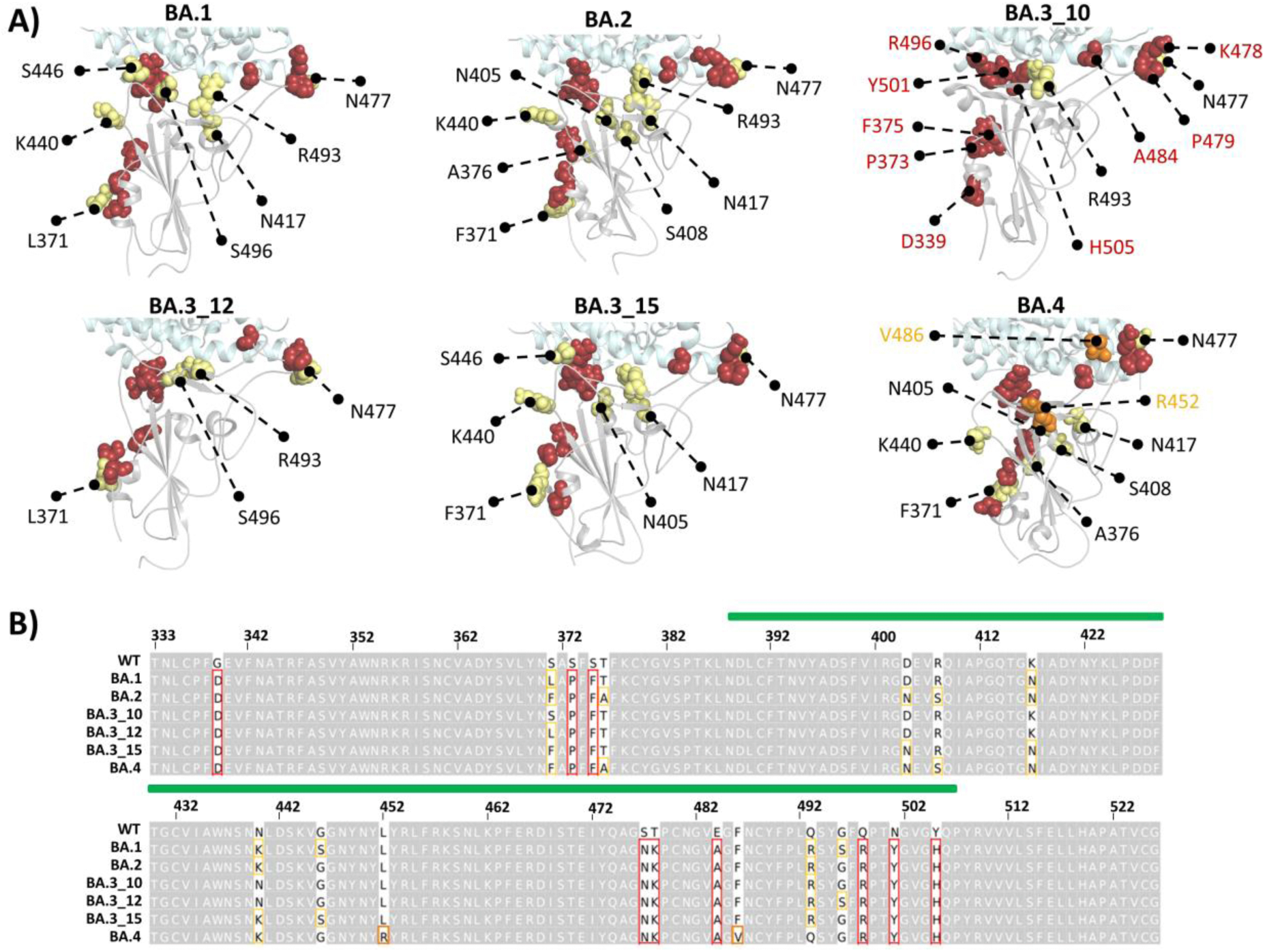
A) Cartoon representation of the RBD structure showing the distribution of the Omicron sub-lineage specific mutations for BA.1, BA.2, BA.3_10, BA.3_12, BA.3_15 and BA.4. The nine mutations common to all sub-lineages are shown as firebrick spheres and annotated in red in BA.3_10 only, whereas the rest of the mutations are presented as yellow spheres. The two unique mutations in BA.4 are shown in orange. B) Alignment of the WT and Omicron sub-lineage RBD protein sequences highlighting the sub-lineage common, unique, and other mutations in red, orange, and yellow, respectively. The RBM region is indicated by a green bar.

To date, several studies identified the effects of Omicron mutations on the RDB domain [12, 68–71] and analyzed the allosteric communications within the S protein [69, 72–74]. For the first time to our knowledge, herein we systematically characterized the collective effects of mutations in each Omicron sub-lineages (BA.1, BA.2, BA.3 and BA.4) both on viral S protein and on human ACE2 protein as complexes and as individual proteins. Trajectory analysis, comparative essential dynamics (ED) calculations and RBD-hACE2 interface residue interaction frequency analysis revealed: 1) a flexible RBM that resulted in conformational variability of the Omicron sub-lineages compared to the WT; 2) highly flexible antigenic hotspots in the RBD which could hinder neutralizing antibody binding; 3) increased residue interactions and interaction frequency between the viral and human proteins in the Omicron variants compared to the WT; 4) significant allosteric effects of mutations in some sub-lineages on ACE2. Furthermore, dynamic residue network (DRN) analysis using our recently developed algorithm [75–78] identified, for the first time, the high centrality communication paths bridging the RBD to the hACE2 core, and the allosteric changes in residue network patterns in both the RBD and hACE2 resulting from collective Omicron sub-lineage mutations.

Taken together, we provide novel insight into the function and evolution of the RBD-hACE2 system which is crucial for drug design for COVID-19 and future viral infections.

## 2. Methods

### 2.1 SARS-CoV-2 Omicron sub-lineage sequence retrieval and structure modeling

Fifty six SARS-COV-2 Omicron sub-lineage sequences were retrieved from Global Initiative on Sharing Avian Influenza Data (GISAID) [79] by searching with the sub-lineage IDs; BA.1, BA.2, BA.3, BA.4 and BA.5 as designated by the PANGO database (https://cov-lineages.org/lineage_list.html). Complete Omicron sub-lineage sequences of African origin, with high coverage and patient status deposited until 24 April 2022 were retrieved from GISAID and submitted to the GISAID in-house tool, CoVsurver [80], which compared them to the SARS-CoV-2 reference sequence: hCoV-19/Wuhan/WIV04/2019 (GISAID ID: EPI_ISL_402124) and identified the sequence specific mutations. RBD specific mutations were extracted via an ad hoc Python script.

The 3D structures of six sub-lineage RBDs in complex with the N-terminal domain of the hACE2 protein were generated using the SARS-COV-2 reference structure (PDB ID: 6M0J) as the template in PyMOL (version 2.5) [81]. All the titratable residues were protonated at a neutral pH of 7.0 using the PROPKA tool from PDB2QR [82] (version 2.1.1) prior to minimization. Please note that, when it is clear from the context, these complexes, and the individual S and hACE2 protein domains of each complex will be referred to by the relevant Omicron sub-lineage name (i.e., BA.1, BA.2, BA.3_10, BA.3_12, BA.3_15 and BA.4).

### 2.2 All atom molecular dynamic (MD) simulations and trajectory analysis

MD simulations using GROMACS [67] v2019.4 were applied to the RBD-ACE2 reference structure (also referred to as wild type, WT) and to the Omicron sub-lineage complexes. Here, *gro* and *top* files were generated from the protonated WT and Omicron sub-lineage systems using the GROMOS54a7 force field, and the structures were placed in a cubic box of 1 nm clearance before being solvated by the single point charge 216 (SPC216) water model. Subsequently, the system charge was neutralized using NaCl ions at 0.15 M concentration. Neutralization was followed by minimization via the steepest descent energy minimization algorithm. An energy step size of 0.01 was used without constraints until a tolerance limit of 1000.0 kJ/mol/nm was reached. Temperature equilibration (NVT ensemble: constant number of particles, volume, and temperature) was achieved using Berendsen temperature coupling at 300 K for 100 ps. Equally, for 100 ps, pressure equilibration (NPT ensemble: constant number of particles, pressure and temperature) was achieved using Parrinello–Rahman barostat [68] at 1 atm and 300K. Thereafter, production runs were performed for 100 ns for each system with a time step of 2 femtoseconds (fs). Under the LINCS algorithm, all bonds were constrained for the equilibration and production runs [69]. Particle Mesh Ewald (PME) electrostatics [70] were used for long-range electrostatic calculations with a Fourier spacing of 0.16 nm. For the short-range Coulomb and van der Waals interactions, a cut-off distance of 1.4 nm was used. All the MD calculations were run at the Centre for High-Performance Computing (CHPC).

Post MD analysis included removal of periodic boundary conditions (PBC) and use of the GROMACS built-in *gmx rms*, *gmx rmsf* and *gmx gyrate* tools to calculate the root mean square deviation (RMSD), root mean square fluctuation (RMSF) and radius of gyration (Rg), respectively. Post MD analysis data were inspected and presented using Seaborn [83], Matplotlib [84], Numpy [85] and pytraj [86] Python packages. Trajectories from MD simulations were viewed using the Visual Molecular Dynamics (VMD) [87] tool. For the WT and each sub-lineage system, the RBD center of mass (COM) in relation to the COM of hACE2 was calculated using the *gmx distance* tool.

### 2.3 Dynamic residues network analysis

DRN calculations average the residue interaction network (RIN) metrics over MD simulations [88, 89]. Here, intra-protein and inter-protein residue networks in the RBD-hACE2 complex were investigated by DRN analysis as applied to the last 20 ns of each MD trajectory for each system. Residues are the nodes in networks and if a connection between two nodes with a Euclidian distance of 6.7 Å of each other exists, it is treated as an edge [89– 91]. The 6.7 Å between residue pairs is a predetermined cut-off distance within the range of ∼6.5-8.5 Å corresponding to the first coordination shell [90]. In a protein network, the residue coordination shell is the range with the highest likelihood of finding residue pairs. Furthermore, while smaller or larger cut-off values generally work in simpler DRN metric calculations, convergence problems arise for larger values for metrics based on shortest path calculations or those that solve for eigenvectors [88].

DRN analysis as established in the MDM-TASK-web [88] includes several centrality metrics, each highlighting a unique node characteristic in the network. Here, five DRN metrics namely, averaged *betweenness centrality* (*BC*), averaged *closeness centrality* (*CC)*, averaged *degree centrality* (*DC*), averaged *eigenvector centrality* (*EC*) and averaged *katz centrality* (*KC*) were calculated (Table 1) for each snapshot of the last 20 ns of the systems’ trajectories using the MDM-TASK-web webserver scripts [88, 89] which are available on GitHub (https://github.com/RUBi-ZA/MD-TASK/tree/mdm-task-web). In the time-averaged form in DRN, *BC* provides a measure of usage frequency of each node by calculating the number of shortest paths passing through a node for a given residue network. Thus, residues with high *BC* values are regarded as functionally and/or structurally significant for that specific network [92]. Averaged *CC* measures the degree of proximity of a node to all the other nodes in the network by computing the average shortest path of a given node to all others. Averaged *DC* is a measure of the connectedness of a node based on the number of unique edges to/from the target node. The more connected a node is to its neighbours, the higher the *DC*. *EC* reflects on the influence of a node in the network based on the centrality of its neighbouring nodes both high scoring and low scoring. Likewise, the *KC* metric indicates the relative influence of a node in a network while considering not only its immediate neighboring nodes but also the neighbors of neighbors. Here DRN was computed for each RBD-hACE2 complex system.

**Table 1:**
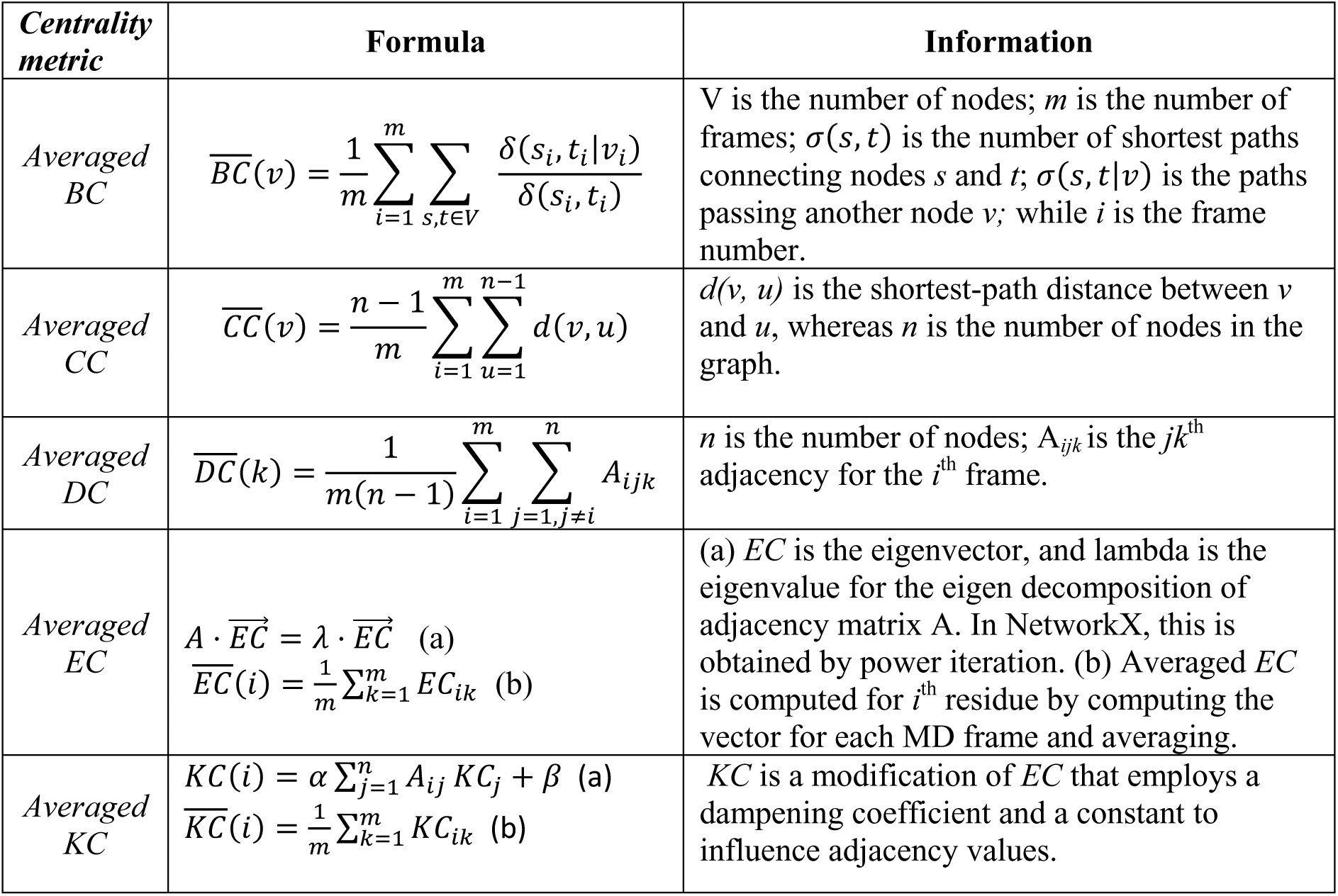
Equations for each DRN centrality calculation [88].

### 2.4 Description of RBD and hACE2 DRN centrality hubs per metric

Centrality is a measure of how central a node is in a protein network and indicates the importance of that residue in communication. Previously, we defined “centrality hubs” as any node that forms part of the set of highest centrality nodes for any given averaged centrality metric [76, 78]. Here, the metric specific DRN centrality hubs were identified using the previously applied analysis algorithm [75, 77, 78]. The algorithm vectorises metric specific results of all the systems in descending order before ranking them and obtaining the hub-threshold value based on a set percentage cut off. Even though DRN was computed for each RBD-hACE2 complex system, the centrality hubs were identified for RBD and hACE2 proteins separately per system due to size difference of the proteins. For that, we used the top 5% for RBD and the top 4% for hACE2 protein as a cut-off value. The established threshold for each case was used to create a binary matrix identifying the hubs and homologous non hub residues as 1 and 0, respectively. The analyzed centrality hub data was rendered as heat maps using Seaborn [83] and Matplotlib [84] Python libraries.

### 2.5 Contact map analysis

Residue contact maps were used to determine the interaction frequency between a given set of residues within a Euclidian distance of 6.7 Å of each other. This cut-off value is used for the reasons explained in Section 2.3. Through contact map analysis, one can identify the gained and lost residue interactions in a network. Interface residues of the RBD in the low energy structure of the reference RBD-hACE2 protein complex that were extracted by comparative essential dynamics (ED) (as described in the next section), were identified using the ROBETTA webserver [93]. This information was used to calculate the contact frequencies over the 100 ns of the MD simulation in each case, and presented as heat maps using the *contact_map.py* and *contact_heatmap.py* scripts (https://github.com/RUBi-ZA/MD-TASK/tree/mdm-task-web) from the MDM-TASK-web [88], respectively. For comparison, a data frame of each residue pair contact frequency per system was created from the *contact_map.py* results and presented as one heat map.

### 2.6 Wild type and Omicron sub-lineage RBD-ACE2 comparative essential dynamics

The most dominant protein motions explored by the Omicron RBD and hACE2 systems were investigated using comparative essential dynamics (ED) [88]. Per system analysis was done by comparing the dynamics of the RBD and hACE2 proteins separately, to that of the WT along principal components (PCs) 1 and 2 using the *compare_essential_dynamics.py* script (https://github.com/RUBi-ZA/MD-TASK/tree/mdm-task-web) from the MDM-TASK-web webserver [88]. The script performed pairwise alignment of each sub-lineage trajectory to that of the WT reference structure via the *Cα* atoms before decomposition of the variance-covariance matrix. Due to increased flexibility, the last three C-terminal residues were excluded from each trajectory. This approach enabled a pair-wise comparison of the prominent motions between the WT and Omicron sub-lineage RBD systems as well as the hACE2 within relevant protein complex. The prominent motions were shown as scatter plots, as described by PC1 and PC2. The scatter plots also indicated the timestamps in picoseconds (ps) for the lowest energy conformations as calculated from 2D kernel density estimates. Furthermore, to enable binding energy computation for near-native low energy complex structures, comparative ED was repeated for the RBD-hACE2 complexes. The low energy structures were extracted using *gmx trjconv tool* and submitted to the HawkDock webserver [94] for binding energy computation.

### 2.7 Dynamic cross-correlation

In dynamic cross-correlation (DCC), we exploit the dynamic nature of protein structures to study their internal movements to decipher the intra-protein and inter-protein interactions and behavior. DCC uses the trajectory and topology files from MD simulations to describe parallel motions of atoms to each other in a protein system. Here, the *calc_correlation.py* script (https://github.com/RUBi-ZA/MD-TASK/tree/mdm-task-web) from MDM-TASK-web [88] as used to rank the degree of the atom correlation in the RBD and hACE2 proteins separately as well as in each system as a whole. The script used the *Cα* atoms from the last 20 ns of each trajectory to rank the internal motions on a scale of -1 to 1, where -1 indicates complete anti-correlation, 1 shows absolute correlation, and 0 means no correlation.

## 3 Results and Discussion

### 3.1 Physicochemical properties of roughly half of the Omicron sub-lineage RBD mutations are not conserved

At the time of the study (April 2022), there were 56 complete sequences for the BA.1, BA.2, BA.3 and BA.4 Omicron sub-lineages from human hosts of African origin in GISAID, with patient status and high coverage (**Table S1**). Unique RBD mutations in the retrieved Omicron sub-lineage sequences were analyzed using the GISAID CoVsurver tool (**Table S2**). Although the WHO lists BA.5 as a new Omicron sub-lineage, no GISAID sequences were retrieved for the sub-lineage under the mentioned search criterion at the time of the study.

To understand the properties and distribution of the RBD mutations,, the sub-lineage specific mutations were mapped to the structure (**Figure 2A**) and the sub-lineage sequences aligned using the Clustal Omega web tool [95] (**Figure 2B**). Here, a minimum of 10 RBD mutations were identified in the BA.3 sub-lineage, which also consisted of sequences with 12 and 15 mutations (referred to from here on as BA.3_10, BA.3_12 and BA.3_15, respectively). The other sub-lineages included 15 mutations in BA.1, 16 mutations in BA.2, and 17 mutations in BA.4. The Omicron sub-lineages shared nine common mutations: G339D, S373P, S375F, S477N, T478K, E484A, Q498R, N501Y and Y505H in the RBD of which, G339D, S375F, S477N and T478K are linked to neutralizing antibody escape [20, 54, 56]. L452R and F486V were unique to the BA.4 sub-lineage. Most Omicron sub-lineages had more RBD mutations than the initial Omicron variant, B.1.1.529 which consisted of 15 RBD mutations, namely G339D, S371L, S373P, S375F, K417N, N440K, G446S, S477N, T478K, E484A, Q493R, G496S, Q498R, N501Y and Y505H [96, 97]. Mutations K417N, G446S, N501Y and Y505H are in residues that form part of the RBD-ACE2 interface and interact with residues in sub-domain I of hACE2 (**Figure 1**).

Thirteen of the twenty-one unique Omicron sub-lineage mutations studied here involved residue substitutions with the same physicochemical properties (**Table S3**). Residues at positions 339, 452, 478, 493 and 498 changed from a non-polar/uncharged residue in the WT to a polar/positively charged residue in the Omicron sub-lineage. The hACE2 interface is predominately negatively charged [98]; consequently, a more positively charged RBD interface would suggest increased RBD-hACE2 electrostatic interaction.

According to the literature, most RBD single mutations reduced the affinity of the S protein for ACE2 but still allowed sufficient levels of expression and ACE2 binding affinity to permit infection, suggesting a high adaption and tolerance to variation in the S protein RBD [99]. Single mutations G339D, N440K, T478K, S477N and N501Y increased the affinity of the RBD for ACE2, while single mutations S375F, K417N, G446S, G496S and Y505H reduced ACE2 binding affinity [99, 100]. S371L, S373P, E484A, Q493R and Q498R were neutral and did not alter the S-ACE2 binding affinity. The E484A mutation did not affect the S affinity for ACE2, but E484K increased ACE2 binding affinity and contributed to immune escape [99, 100]. The N501Y mutation implicated in strengthening RBD-hACE2 binding [89] involves substituting an asparagine with a tyrosine residue. While both residues are polar and capable of hydrogen bonding, the tyrosine ring would potentially alter the interface interactions through pi-stacking and pi-cation interactions. In the BA.4 sub-lineage, the unique L452R and F486V mutations increased and decreased the affinity for ACE2, respectively [99]. Additionally, residues at positions 446, 496 and 493 reverted to the WT sequence compared to the previous lineages (i.e., Gly446, Gln493, and Gly496). While the Q493R mutation was neutral, the wild type G446 and G496 residues in BA.4 would increase S-ACE2 affinity compared to the serine substitutions observed in the previous sub-lineages. It is interesting to note that single mutation analyses did not always result in increased affinity i.e., a combination of both increased and decreased affinity mutations were observed. In our recent review article, we indicated that not all non-catalytic site mutations have an allosteric effect on the function of the protein unless combined with other mutations; which we refer to as neutral mutations, [76]. We further suggested that collective analysis of mutations is needed to provide insight into mechanisms [76], similar to allosteric polymorphism, in which mutation of several critical positions in the protein sequence allosterically disrupt the protein function [101]. This may suggest evolution towards an ‘induced fit’ in the Omicron sub-lineages in which affinity has to be lost via mutations in some residues in order to support the gain of affinity at another site. Considering this, hereafter each sub-lineage mutations were analyzed collectively.

### 3.2 Omicron sub-lineage RBD mutations collectively influence the RBD-hACE2 complex dynamics

The reference RBD-hACE2 protein complex (6M0J) and the modeled Omicron sub-lineage RBD structures complexed with the N-terminal domain of the hACE2 reference protein were subjected to 100 ns all-atom MD simulations, and further evaluated through trajectory analysis using RMSD, RMSF, Rg and comparative ED.

The RBD-hACE2 reference (WT) RMSD results from the quality control duplicate runs showed agreement in system equilibration over the 100 ns simulation time (**Figure S1A**). As shown in the RMSD line plots in **Figure S1B**, most of the systems behaved like the WT, except BA.3_12. RBD RMSD violin plots (**Figure 3A**) showed that, except for BA.2, all other RBD proteins including the WT had at least a bimodal RMSD distribution. BA.3 and BA.4 proteins experienced high structural variations compared to the WT based on the median RMSD. BA.2 was particularly interesting with its unimodal behavior. We also examined the hACE2 RMSDs violin plots to determine if the RBD mutations have any effect on hACE2 within each protein complex. The unimodal behavior of the WT hACE2 was maintained in all except BA.3_15 (**Figure 3B**). Overall, the RBD proteins with sub-lineage mutations experienced notable structural variations, which did not directly affect hACE2 protein in each of the systems, except for BA.3_15.

**Figure 3:**
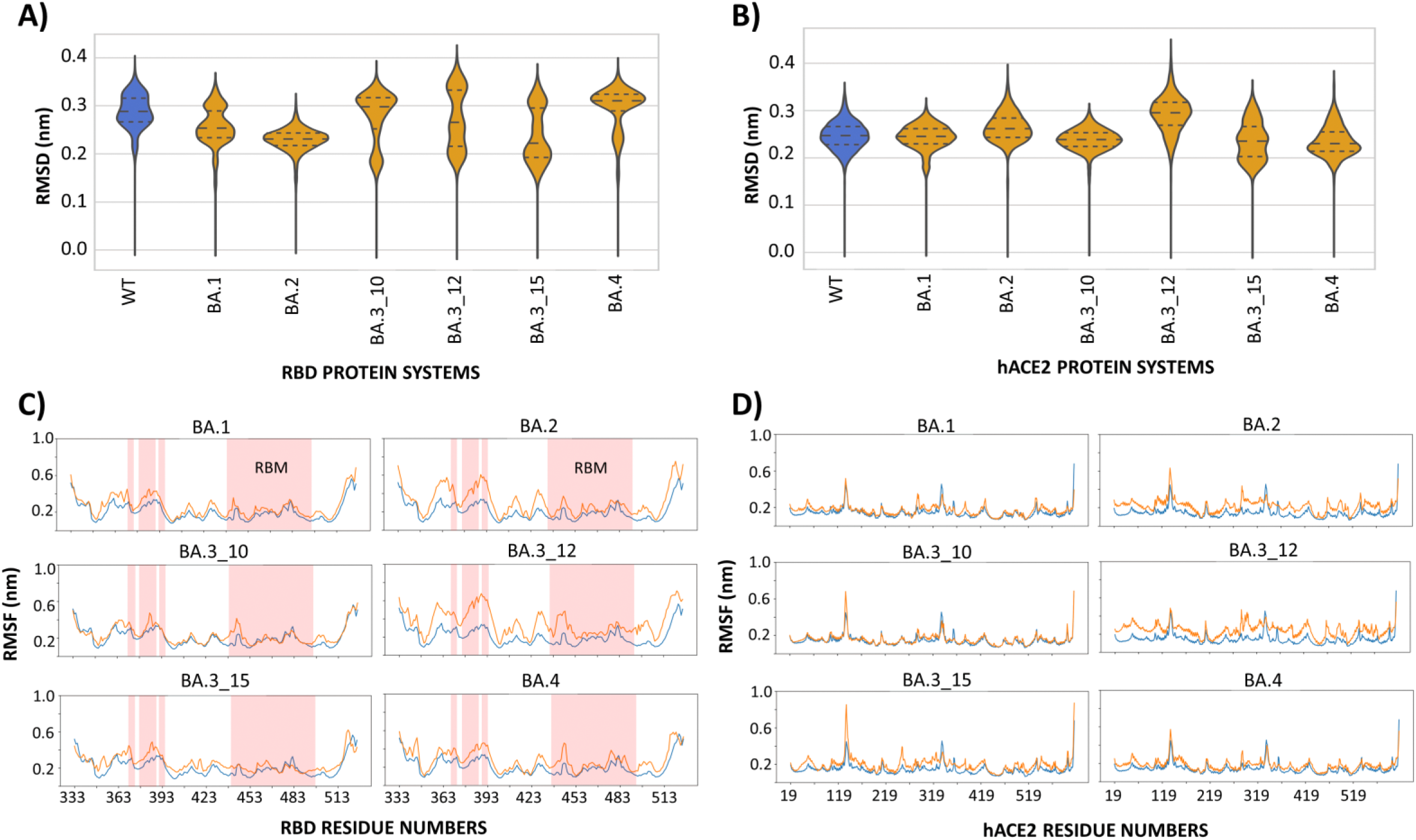
RMSD violin plot distribution of A) RBD and B) hACE2 within each complex. C) and D) show the comparative RMSF distribution between of the WT (blue) and mutant system (orange) for the RBD and hACE2 proteins, respectively. The x and y-axes show the RMSF values and residue positions, respectively. RBM region, 438-508 and antigenic sites, 370, 375-386, 390 and 444-456 are marked on RBD RMSF line plots.

RMSF calculations for the RBDs (**Figure 3C**) indicated increased residue fluctuation across the sub-lineage systems compared to the WT. Considerable residue flexibility was noted around positions 348 to 393 and 423 to 453, specifically in BA.2 and BA.3_12 in contrast to the WT. RDB positions 370, 375-386, 390, 444-456 are identified as antigenic sites recognized by neutralizing antibodies in the S RBD protein [85]. Increased residue fluctuations around the antigenic sites in the RBD may be a strategy for neutralizing antibody escape by preventing stable binding and hence reducing maturation of high-affinity antibody paratopes [102]. Structural flexibility at antigenic sites may also prevent appropriate immune recognition and reduce antibody production [103]. It is also likely that the increased RBD dynamics could expose the S RBD to more interactions with the host receptor, hACE2, thereby improving binding [12]. This is further discussed in **Section 3.6**. From the hACE2 perspective, BA.2, BA.3_12 and BA.3_15 had greater residue flexibility compared to the WT (**Figure 3D**).

The Rg analysis showed nominal differences between the Omicron sub-lineage systems and the WT. However, BA.4 had the highest Rg in the RBD (**Figure S1C**), whereas BA.2 and BA.3_12 had a higher hACE2 Rg than the WT (**Figure S1D**).

### 3.3 Comparative essential dynamics calculations revealed more conformational space in Omicron sub-lineage systems compared to the WT

We further investigated conformational evolution of the individual proteins in the complexes using a comparative ED approach. In ED, the dominant protein motions are determined by decomposing the variance-covariance matrix obtained from the *Cα* or *Cβ* atom positional changes during MD simulations. The comparative ED approach from MDM-TASK-web [73] further describes the dominant motions of two or more systems within the same Eigen subspace. In our case, we compared each protein to the reference protein.

From **Figure S2A**, PC1 and PC2 accounted for ∼50% of the RBD motions in all the systems. All Omicron systems sampled more conformational space than the WT. In BA.1, PC1 and PC2 explained ∼44% of the conformational variance with most variation compared to the WT along PC2. Similarly, BA.2, BA.3_10, BA.3_12 and BA.3_15 experienced more RBD conformational variation along PC2 compared to the WT. PC1 and PC2 accounted for ∼57, 52, 51 and 60% of the motions in these systems, respectively. Compared to the WT, BA.4 experienced more RBD conformational diversity along both PC1 and PC2, which accounted for ∼48% of the observed conformational variance.

From the hACE2 perspective, BA.2, BA.3_12 and BA.3_15 experienced a more dispersed conformational space along both PC1 and PC2 compared to the WT (**Figure S2B**). Here, PC1 and PC2 collectively described ∼52, 61 and 56% of the observed variance, respectively. Compared to the WT, the most conformational variance in BA.1 and BA.3_10 was along PC2. The collective PC1 and PC2 variance was above 50% in both systems. BA.4 had more slightly distributed conformational space along PC1 and PC2 with a total variance of ∼50% compared to the WT.

### 3.4 Binding of Omicron sub-lineage RBDs allosterically affected the hACE2 substrate binding pocket conformation

Binding of the SARS-CoV-2 RBD to ACE2 affects the carboxypeptidase activity of the enzyme, with RBD binding being sufficient to increase ACE2 catalytic activity and substrate affinity against model peptides substrates, caspase-1 and Bradykinin analog [104–106]. The RBD of the SARS-CoV-2 S protein binds to subdomain I in the N-terminal domain of the hACE2, the same sub-domain which undergoes a substantial hinge-like movement upon ligand binding to enclose the substrate in the central protease cavity [35]. In contrast to this conformational change, the high affinity binding SARS-CoV-2 RBD (and not the SARS-CoV RBD which bound ACE2 with lower affinity) induced conformational changed in sub-domain II of ACE2, resulting in closing of the active site and tighter binding of substrate peptides [106]. We therefore sought to investigate the binding effect of RBD on the hACE2 substrate binding pocket conformation. We defined the substrate binding pocket in two different ways: 1) the entire groove between the sub-domains I and II; 2) the pocket defined by the active site residues, Phe274, Leu278, His345, Asp368, Thr371, Glu375, His378, Glu402, His505, Tyr510, Arg514 and Tyr515.

Comparative ED calculations as applied to the entire groove region showed more conformational diversity in the hACE2 groove cavity in the Omicron sub-lineages than the WT (**Figure S3A**). Conformational diversity was particularly pronounced in BA.2, which had the most variation along PC1 and PC2, accounting for ∼56% of the dominant motions, and BA.3_12 along PC1 and PC2 with a total variance of ∼67%. We also analyzed the groove cavity via Rg calculations, and the results indicated that BA.2 and BA.3_10 had slightly less pocket compaction compared to the WT (**Figure S3B**). The conformational changes in the entire groove region observed via ED and Rg could imply that the RBD binding affects the hACE2 substrate pocket dynamics in the Omicron sub-lineages. RBD-induced closing of sub-domain II was linked to increased substrate affinity [106].

On the contrary, COM calculations as defined by active site residues, showed minimal changes in the distance between the active site residues in sub-domain I (His345, Asp368, Thr371, Glu375, His378, Glu402) and sub-domain II (Phe274, Leu278, His505, Tyr510, Arg514, Tyr515) for the Omicron sub-lineages compared to the WT (**Figure S4A**). This was further supported by Rg calculations of the active site pocket which showed a small range in the gyration radius of 1.00 - 1.4 nm for all the systems (**Figure S4B**). Taken together, these results indicate that binding of Omicron sub-lineage RBDs has noticeable effects on entire groove region of ACE2, but the change to the catalytic site is minor. It is worth noting that the distance between the active site residues and the substrate binding pocket are critical for ACE2 catalytic activity [31].

From a global perspective, the RBD Omicron sub-lineage mutations affect the conformational behavior of the RBD itself and the hACE2 protein at different levels, as demonstrated by the RMSD, Rg and especially comparative ED calculations. Dehury et al. previously showed an inward motion of the RBD towards the hACE2 in the Y489A and Y505A RBD mutant systems [107]. To further understand the mutational effect on inter-protein dynamics, we employed COM distance and dynamic cross-correlation calculations in the next section.

### 3.5 Anti-correlated RBD-hACE2 motions and increased inter-protein interaction space was observed in some Omicron sub-lineage systems

We studied atomic correlations within the RBD and hACE2 proteins, as well as between the RBD-hACE2 complexes via dynamic cross-correlation (DCC) analysis. In the RBD-hACE2 complex, correlated atomic motions between the two proteins were noted in the reference structure compared to Omicron sub-lineage complexes BA.1, BA.3_10, BA.3_15, and BA.4 (**Figure S5A**). In the sub-lineage structures, the anti-correlated motions were mainly observed between the RBD and hACE2 atoms, implying opposing movements between the proteins. This observation was concordant with the Omicron sub-lineage RBD-hACE2 COM distance analysis, which also showed a marginally higher COM distance in the sub-lineages compared to the WT (**Figure S6**). Furthermore, binding energy calculations using low energy structures, extracted according to comparative ED results, showed that the WT had a lower protein binding energy (-80.6 kcal/mol) compared to the majority of the Omicron sub-lineages: BA.1 (-46.45 kcal/mol), BA.2 (-55.94 kcal/mol), BA.3_10 (-82.72 kcal/mol), BA.3_12 (-82.72 kcal/mol), BA.3_15 (-93.78 kcal/mol) and BA.4 (-74.4 kcal/mol).

We further focused on individual proteins in the complexes and calculated DCCs for each protein. RBD focused intra-protein DCC analysis identified atomic correlations in the WT, BA.3_10, BA.3_15 and BA.4 Omicron sub-lineages (**Figure S5B**). Intra-protein anti-correlated atomic motions were noted in BA.1, BA.2 and BA.3_12, particularly between residues 440 and 508. This region encompasses the receptor binding motif (RBM) loop region responsible for hACE2 binding [8] (**Figure 1**). The increased RBM flexibility in the Omicron sub-lineages was earlier noted in the RMSF calculations (section 3.2) (Figure 3C). Interestingly, BA.1, BA.2 and BA.3_12 also experienced anti-correlated atomic motions in the hACE2 especially at regions 119-315 and 419-519 which mainly form part of sub-domain II (**Figure 5C**). This implied that RBD dynamics, especially in the RBM, influence motions of regions of ACE2 distinct from the S binding site. This is consistent with observations of conformational changes in sub-domain II caused by S2 RBD binding [106], which contrast with sub-domain I conformational changes induced by ligand binding during which sub-domain II was largely static [35]. The global changes in protein-protein dynamics due to sub-lineage mutations suggest that the RBD mutations influence atomic interactions and possibly communication patterns. Our previous studies showed changes in communication and allosteric paths resulting from SNPs in several proteins [77, 79, 80]. This approach was applied to the RBD-hACE2 systems, as discussed in the next section.

**Figure 4:**
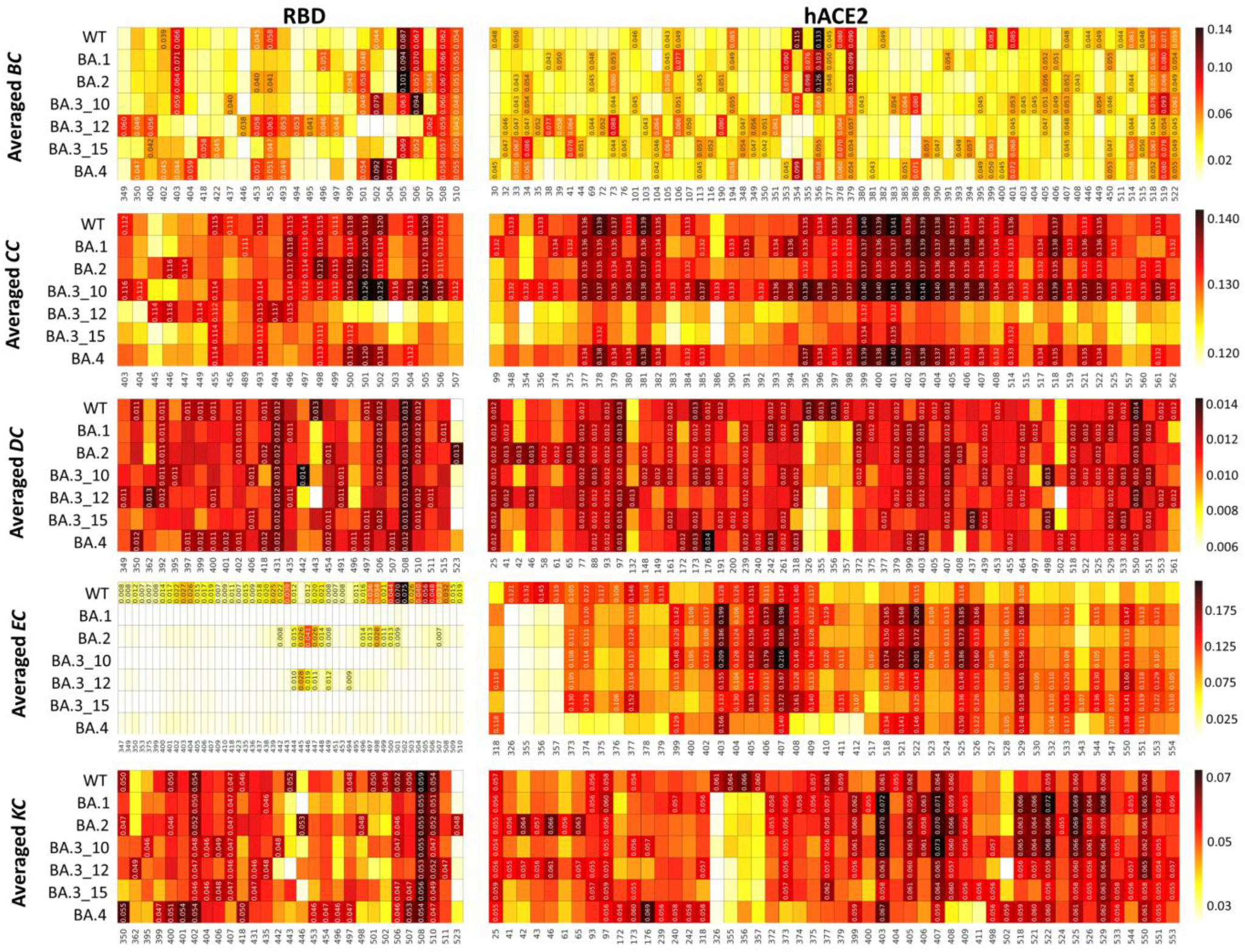
Heat map representation of hubs per DRN metric as calculated using the global top 5% for the RBD and 4% for the hACE2 proteins. Hub residues are annotated with centrality values whereas their homologous residues from other systems are not. Hub residues are shown on the x-axis and the protein systems on the y-axis. The color scale from white through red to black indicates the residue centrality values.

**Figure 5:**
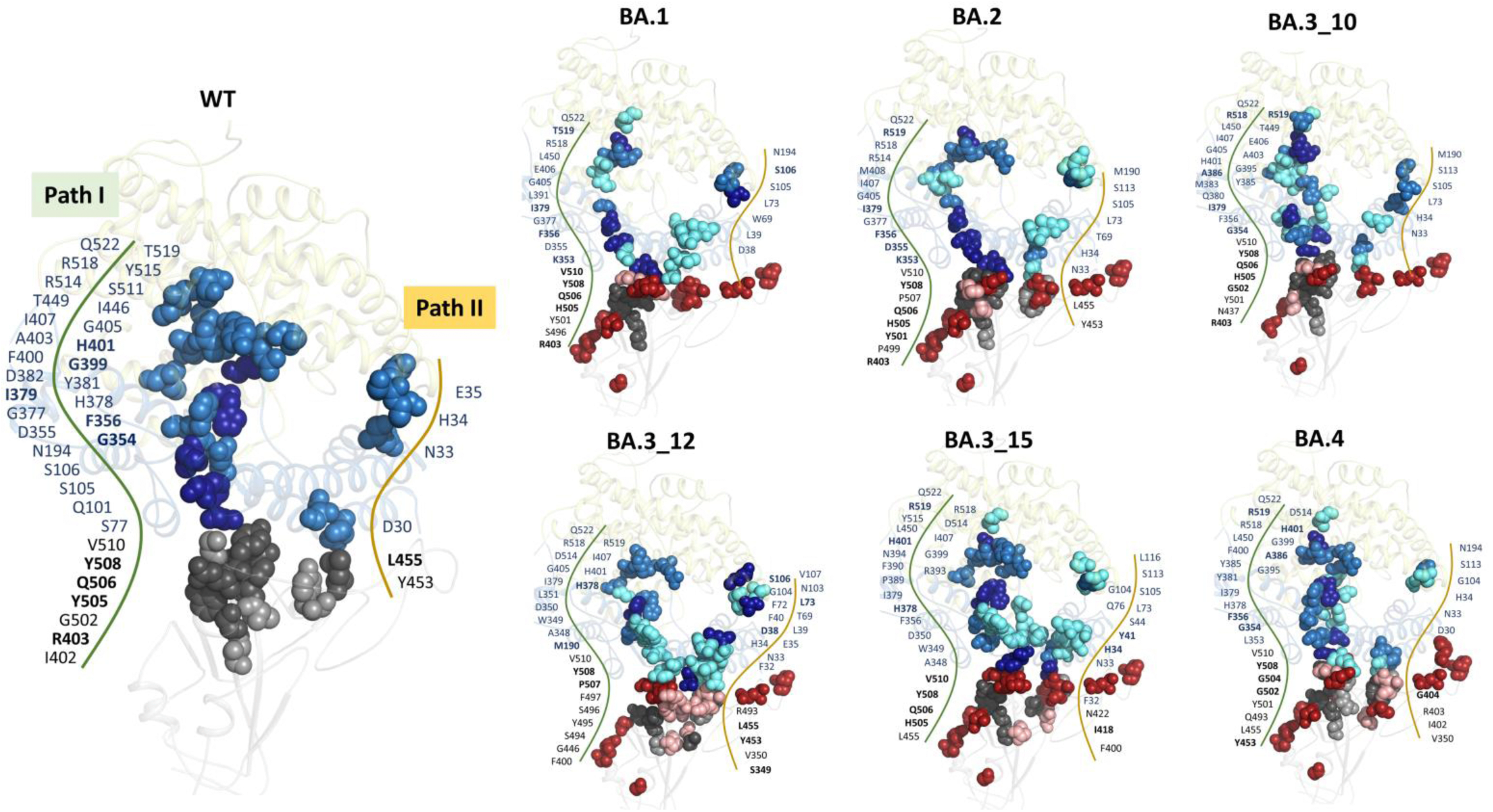
Cartoon representation of the RBD-hACE2 structures showing the distribution of the global top 5% and 4% *BC* hubs in the RBD and hACE2, respectively for the WT and Omicron sub-lineages. The RBD is shown in grey and hACE2 sub-domains I and II as sky-blue and pale-yellow, respectively. WT hubs are shown as sky-blue spheres (hACE2) and grey spheres (RBD). The same colors are used for *BC* hubs common to the WT and Omicron sub-lineages. *BC* hubs unique to the Omicron sub-lineages (Δ hubs: sub-lineage hubs – WT hubs) are shown as aquamarine spheres (hACE2) and boron spheres (RBD). The five *BC* hubs with the highest centrality values in the RBD and the hACE2 are shown as dark grey and dark blue spheres, respectively, and annotated in bold. The sub-lineage specific mutation positions are shown as firebrick spheres.

### 3.5 Dynamic residue network analysis was performed for five different metrics

Protein molecules exist as a network of amino acids whose atomistic contacts facilitate intra-protein, inter-protein communications and ligand/receptor binding and interaction [108, 109]. The networked nature of protein residues can be represented as nodes and the pairwise connections between residues as edges where relationships can be studied using graph theory [88, 89]. Consequently, changes in the amino acid composition due to mutations which affect both intra-protein and inter-protein interaction patterns can be investigated through network analysis.

Here, like in our previous studies [75, 77, 110] we employed five DRN analysis metrics; averaged *betweenness centrality (BC)*, averaged *closeness centrality (CC)*, averaged *degree centrality* (*DC)*, averaged *eigenvector centrality (EC)* and averaged *katz centrality* (*KC)* to explore the communication network differences between the WT and the Omicron sub-lineage RBD-hACE2 protein complexes. The centrality hubs for each metric were identified using a global cutoff of top 5% for the RBD and 4% for the hACE2 proteins. Results were presented as heat maps indicating system specific hub residues per metric and their corresponding homologous residues in the other systems (**Figure 4**).

#### 3.5.1 Betweenness centrality identified two distinct allosteric communication paths between RBD and hACE2 that progressively evolved through the sub-lineages

The *BC* metric assigns centrality based on the usage frequency of a residue in the shortest paths calculated between all possible residue pairs within the given network [111, 112]. We assume that within a protein (or protein complex) the communication goes through the shortest paths, hence residues with high *BC* values within these paths are regarded as functionally important, especially in the control of information flow [91].

According to **Figure 4**, Tyr508 and Val510 hub residues were unaltered from the reference protein (WT) in the presence of Omicron sub-lineage mutations for the averaged *BC* calculations; thus, they are the *persistent hubs*. We previously introduced the term “*persistent hubs”* to define the hubs that remain unchanged within a set of comparative systems, and thus *persistent hubs* are the indication of the functional importance of these residues [78]. Tyr508 is located just outside the RBM region at the N-terminus end of the β7. Tyr508 is involved in the RBD-hACE2 interaction [113, 114], binding and interaction with camel nanobodies for RBD neutralization [115], standard drugs [116, 117], inhibitory peptides [118], metal complexes [119] and natural inhibitory compounds [120]. Val510 binds a number of natural bioactive compounds with potential antiviral properties [116, 121, 122]. Mutations in either Tyr508 or Val510 reduce RBD-ACE2 binding affinity [99].

In the hACE2, *BC* hub residues Ile379, Arg518, Thr519 and Gln522 were *persistent* across all the systems (**Figure 4**). The other *BC* hubs present in at least five systems included residues at positions Asn33, His34, Leu73, Phe356, His401, Ile407 and Arg514. Residues Asn33 and His34 are part of the hACE2 sub-domain I helix which is responsible for the majority of RBM interactions, while residue Phe356 lies within the unstructured part of the protein (351-357) that forms additional RBM contacts [6]. His401 is a zinc coordinating residue in hACE2 [35].

Interestingly, we observed two distinct communication paths that bridge the two protein cores by mapping of the WT and sub-lineage RBD and hACE2 averaged *BC* hub residues onto the respective 3D structures (**Figure 5**). In the WT, there was a main network of high centrality hubs (Path I) connecting the two proteins involving RBD and hACE2 residues as indicated in the **Figure 5**. The other high centrality network path (Path II) was much shorter, but still bridged two proteins. RBD residues for Path II are at positions Tyr453 and Leu455; and hACE2 residue positions are 30, Asn33, His34, Gln101, Ser105, Ser106 and Asn194. We further highlighted the top five *BC* hubs of the RBD and hACE2 proteins with the highest centrality values in dark grey and dark blue color in the **Figure 5**. One of these highest centrality hubs of the WT RBD, Tyr505, was mutated to His in all Omicron sub-lineages that we studied, yet remained as hub in only BA.1, BA.2. BA.3_10 and BA.3_15. This mutated residue lost its centrality (hence functional importance) in BA.4. In WT, Tyr505 forms a hydrogen bond with Glu37 and contact interactions with Lys353 and Arg393 of hACE2 [123]. His retains the ability to form hydrogen bonds, and the introduction of a positive charge might increase electrostatic interactions with the predominantly negatively charged ACE2. However, the single mutation of Tyr505 to His reduced affinity of the RBD for ACE2 [99], although in our analysis the Y505H mutation leads to increased binding interface contacts in all mutant protein complexes (discussed further in Section 3.6). This further emphasizes the importance of analyzing the sub-lineage mutations collectively [76].

Other important S protein RBD residues that had *BC* hub status unique to the Omicron sub-lineages included Ser496 in BA.1, Pro499 in BA.2 and Arg493 in BA.3_12 and BA.4. The high centrality observed in the specific sub-lineages, especially for the residues involved in RBD-hACE2 interaction, highlights the importance of these residues in binding and inter-protein interactions of the Omicron sub-lineages.

We also observed compelling changes in these two paths in the Omicron sub-lineages (**Figure 5**). In BA.1, the RBD mutations resulted in loss of hub residues Glu402, Tyr453, Lue455 and Ser502. However, a compensatory gain in centrality was noted involving RBD interface residues Thr496 and Thr501. In fact, N501Y mutation, associated with strengthening the RBD-hACE2 inter-protein interactions [124–126], gained *BC* hub status in BA.2, BA.3_10 and BA.4 too. Similar to BA.1, BA.3_10 and BA.3_15 showed loss of *BC* hubs due to Omicron RBD mutations.. However, in both cases, we identified compensatory gains of centrality hubs both at the interface and core regions of the RBD (**Figure 5**). Omicron sub-lineages, BA.2, BA.3_12 and BA.4 with 15, 12 and 17 RBD mutations, respectively, presented with more RBD *BC* hubs compared to the WT. In these systems, the gain in *BC* hubs was mainly at the interface region involving residues Pro499 and Thr501 in BA.2, Gly446, Arg493, Ser494 and Ser496 in BA.3_12, and Arg493, and Tyr501 in BA.4. Furthermore, there was clustering of *BC* hubs around the hACE2 interface in BA.1, BA.3_10, BA.3_12 and BA.3_15 which implies that the changes in the RDB in turn affect the communication patterns in hACE2 too. More so, the BA.3 sub-lineage had more *BC* hubs (BA.3_10: 25 hubs; BA.3_12: 28 hubs; BA.3_15: 30 hubs) in the hACE2 compared to the WT (23 hubs). In these sub-lineages, an enhanced *BC* Path I bridging the two proteins, was observed involving newly acquired hubs in the hACE2.

Collectively, for the first time, we showed 1) two separate allosteric communication paths between the S RBD and hACE2 (Path I and II) formed via averaged *BC* centrality hubs; 2) changes in the residue network patterns of these two paths in the Omicron sub-lineages highlighting the compensatory gains in *BC* hubs at the RBD interface to maintain cross communication with the receptor; and 3) that the RBD mutations not only influence the communication patterns in the RBD but also enhance the communication path in hACE2 of some sub-lineage systems though residue gain in centrality, especially in the interface area. Overall, the *BC* hubs of the RBD and hACE2 complex suggest an evolutionary progression of the Omicron sub-lineages towards establishment of stronger and more efficient communication paths between the viral S protein and the human ACE2 receptor.

#### 3.5.2 Closeness centrality hubs of sub-lineages supported an evolutionary progression

Previously we showed that interface and/or protein core residues which are proximal to all other residues in the network have high *closeness centrality* (*CC*) values [77, 78, 127]. Here, *CC* calculations again identified RBD interface residues adjoining the hACE2 as key hubs. There were no *persistent hubs* in the RBD, mainly due to the dramatic hub changes observed in BA.3_12 and BA.3_15 sub-lineages. However, residues Leu455, Gln493, Gly496, Gln498, Thr500, Asn501 and Gly502 located at the RBD interface and part of the RBD-hACE2 interaction [6, 30, 128, 129], had high centrality in at least five of the seven systems (**Figure 4**). ln the hACE2, residues Gly399 and His401 were identified as *persistent hubs* across all the systems (**Figure 4**). These residues form part of the catalytic pocket which lies between sub-domains I and II of the N terminal domain of ACE2 [35].

In our previous work, we showed a correlation between the increased COM distance of proteins within a protein complex and reduced number of *CC* hubs. Here, we observed a similar trend for BA.3_12 with an increased COM distance compared to the reference structure (**Figure S6**) and a highly reduced number of *CC* hubs compared to all other systems (**Figure 4 and 6**). As earlier noted from the RMSF results, the hACE2 of the BA.3 systems, especially BA.3_12, showed increased residue flexibility compared to the WT. Since *CC* assigns centrality based on residue proximity to the neighbors, the increase in residue flexibility in the Omicron sub-lineages as well as increased COM distance could explain the fewer number of *CC* hubs, especially in BA.3_12. BA3.15 had also fewer hubs compared to WT, BA.1, BA.2 and BA.4. Interestingly, the opposite behavior was observed for BA.3_10. In this case, the RBD and ACE2 were closer to each other compared to the WT protein complex (**Figure S6**), and we observed the highest number of *CC* RBD and hACE2 hubs of all the systems. These three intermediate BA.3 sub-lineages may represent progressive trial-and-error based evolution by the virus to optimize binding of the RBD to hACE2; while mutations in BA.3_10 make the complex closer and more rigid, the BA.3-12 and BA3.15 mutations weaken the RBD-hACE2 interactions compared to other variants [60, 130].

**Figure 6:**
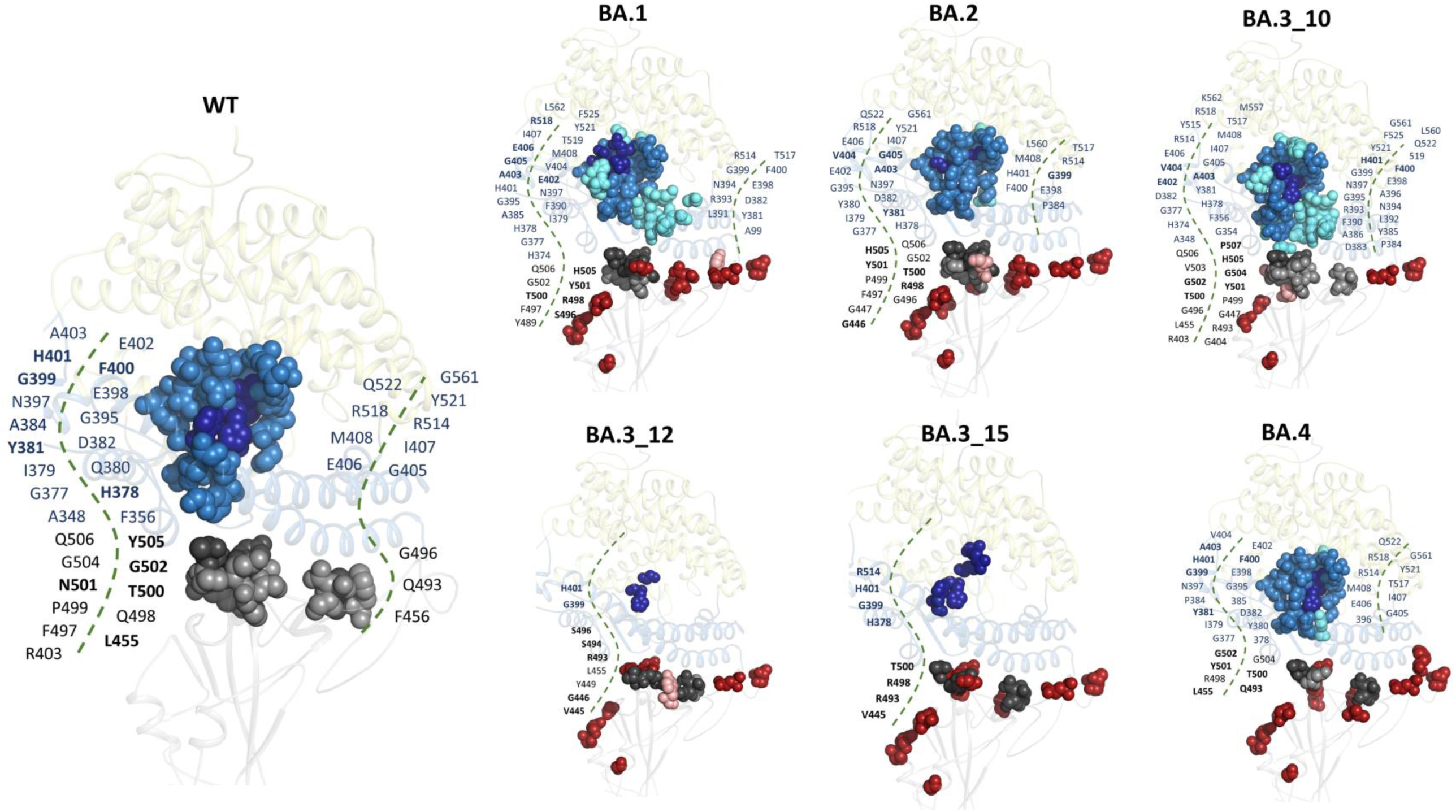
Cartoon representation of the RBD-hACE2 structures showing the distribution of the global top 5% and 4% *CC* hubs in the RBD and hACE2, respectively for the WT and Omicron sub-lineages. WT hubs are shown as sky blue spheres (hACE2) and grey spheres (RBD). The same color is used for *BC* hubs common to the WT and Omicron sub-lineages. *BC* hubs unique to the Omicron sub-lineages (Δ hubs: sub-lineage hubs – WT hubs) are shown as boron spheres whereas mutation positions are shown as firebrick spheres. The five highest centrality residues in RBD and hACE2 are shown as dark grey and dark blue spheres, respectively, and annotated in bold.

To further understand the relationship between the RBD mutations and *CC* hub distribution, a closer evaluation of the evolution of the *CC* hubs with the number of mutations was done (**Figure 6**). Here, we focused on systems with differing numbers of RBD mutations to better explain the relationship. In BA.2, introduction of the extra D405N and R408S mutations absent in BA.1 resulted in the loss of centrality/hub status of residue Tyr489 in BA.2. Tyr489 is key in RBD-hACE2 interactions where it forms hydrogen bonds with Tyr83 [60]. Evidently, from RMSF calculations, a higher flexibility was noted at position 489 in BA.2 (0.2045 nm) compared to BA.1 (0.1794 nm). Additionally, the G446S substitution in BA.1 resulted in loss of *CC* hub status at this position in the sub-lineage compared to BA.2 which lacks this mutation. When comparing BA.2 with BA.3_10, an increase in *CC* hubs is noted in BA3_10, which has fewer mutations (10). BA.3_10, gained *CC* hubs at positions Arg403, Gly404, Leu455, Arg493 and Val503 compared to BA.2. Additionally, mutation Q493R resulted in loss of the *CC* hub at this position in BA.2. Comparison of the BA.3 mutation combinations (10, 12 and 15) clearly illustrated the depreciating effect of the RBD mutations on the protein centrality. Here, progression from 10 RBD mutations in BA.3_10 to 12 mutations in BA.3_12 resulted in a loss of hubs status for residues Arg403, Gly404, Phe497, Gln498, Pro499, Thr500, Asn501, Gly502, Val503, Gly504, Tyr505, Gln506 and Pro507, the majority of which interact with hACE2. Similar observations were made between BA.3_12 and BA.3_15 where, residues, Val445, Gly446, Tyr449, Ser494 and Gly496 lost hub status in BA.3_15. Interestingly, stabilization of *CC* hubs at positions Gly502 and Gly504 was noted between BA.3_15 and BA.4 sub-lineages which differ by 2-mutations. The Omicron sub-lineage BA.4 gained more *CC* hubs at the RBD interface, namely Tyr501, Gly502 and Gly504 compared to BA.3_15, despite having two extra mutations. Here the N501Y mutation resulted in high *CC* at this position as in the WT. More so, there was a reversion to a high number of *CC* hubs in the hACE2 of BA.4 compared to BA.3_15, characterized by new *CC* hubs at Tyr385 and Thr517, like the WT and earlier Omicron sub-lineages. Tyr385 and Thr517 are located in the active site cleft [35].

Together, the *CC* analysis demonstrates how the progressive evolution of Omicron sub-lineages affects the centrality of the S RBD which in turn affects the residue dynamics and interaction with the receptor hACE2.

#### 3.5.3 Allosteric communication path formed by the WT eigen centrality hubs is interrupted in the Omicron sub-lineage RBD-hACE2 complexes

From the global top 5% *EC* calculations, highly influential RBD *EC* hub residues were identified exclusively in the WT, BA.2 and BA.3_12 systems (**Figure 4**). In BA.2, RBD *EC* hubs included residues at positions 422, 444-449, 496-501 and 507; whereas in BA.3_12 *EC* hubs were at positions 444-447, 449 and 494. No RBD hubs were observed in BA.1, BA.3_10, BA.3_15 and BA.4 systems.

Interestingly, structural mapping of *EC* hubs in the WT showed a definite allosteric communication path of *EC* hubs from the RBD core, traversing the RBM of the S protein and the N-terminal domain of the hACE2, to the zinc binding site in the hACE2 (**Figure 7**). In the WT, the *EC* hubs constituted part of the β2, β3, β4 and β7 strands of the RBD core and part of the RBM involving residues at positions 438, 439, 442-447, 449, 451, 453 and 494-506. Residues Ala403, Ile436, Leu444, Thr449, Cys498, Asn501 and His505, which are documented epitopes for neutralizing antibody binding [57, 131, 132], had hub status in the WT which was lost in majority of the Omicron sub-lineages (BA.1, BA3_10, BA3_15 and BA.4). Recently, we showed a relationship between a dynamically stable C-terminal domain of the KatG protein and a high concentration of *EC* hubs in the domain, implying that highly influential residues are associated with stable regions [75]. Likewise, here, the Omicron sub-lineages experienced higher RBD residue fluctuation compared to the WT which could explain the scarcity of *EC* hubs. Furthermore, the loss of *EC* hub status especially in the RBM and RBD neutralizing antibody epitopes of the Omicron sub-lineages signifies a loss of residue influence/centrality at these positions as a potential antibody escape mechanism.

**Figure 7:**
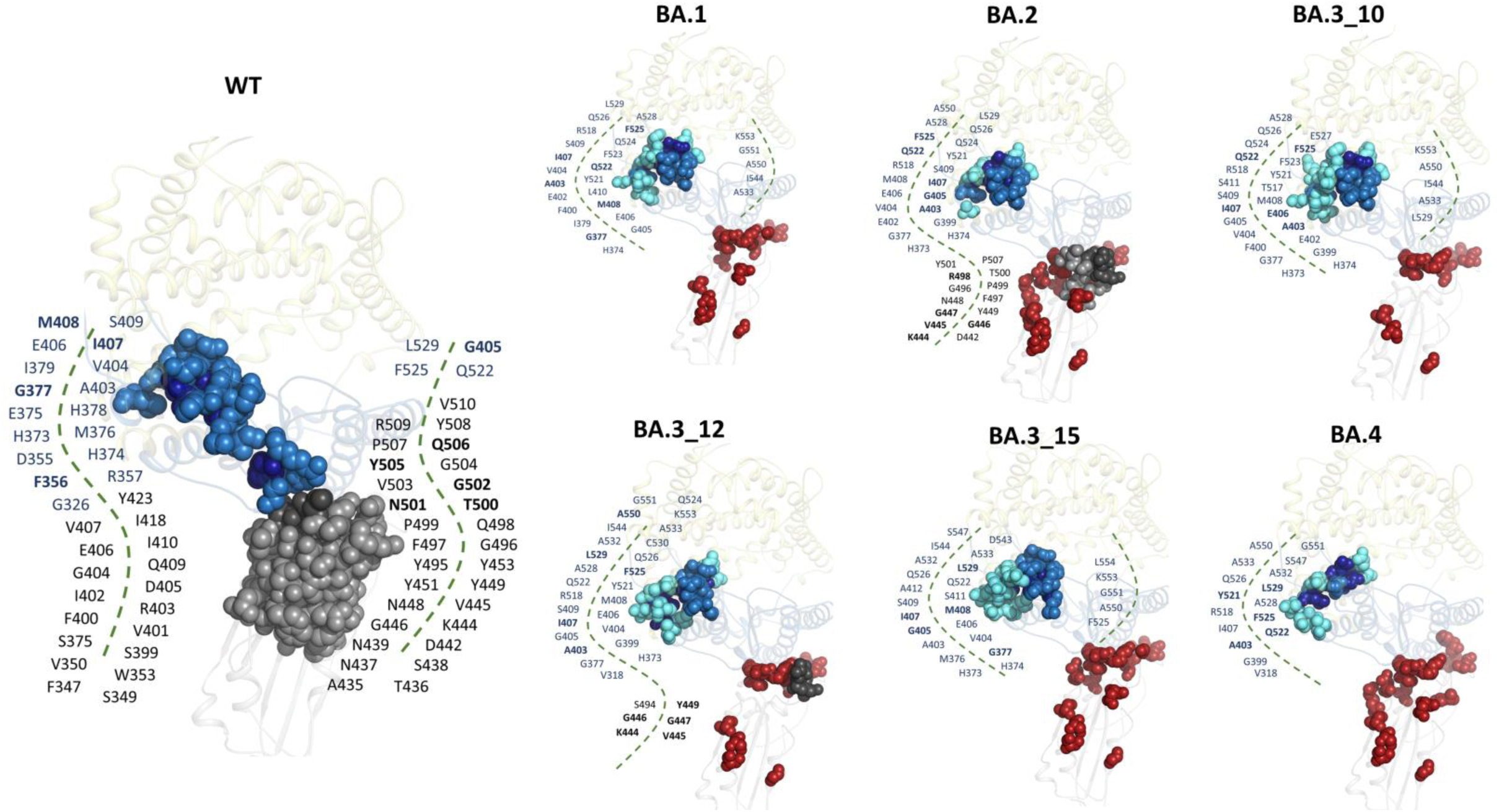
Cartoon representation of the RBD-hACE2 structures showing the distribution of global top 5% and 4% *EC* hubs in the RBD and hACE2, respectively for the WT and Omicron sub-lineages. WT hubs are shown as sky-blue spheres (hACE2) and grey spheres (RBD). The same colors are used for *EC* hubs common to the WT and Omicron sub-lineages. *EC* hubs unique to the sub-lineages (Δ hubs: sub-lineage hubs – WT hubs) are shown as aquamarine spheres (hACE2) and boron spheres (RBD). The five highest centrality residues in RBD and hACE2 are shown as dark grey and dark blue spheres, respectively, and annotated in bold. Omicron sub-lineage specific mutation positions are shown as firebrick spheres.

This communication path also included hACE2 residue positions 326, 355-357, 375, 378 and 379 only in the WT. These residues are part of sub-domain I and include the zinc coordinating residue His378. Furthermore, hACE2 residues in sub-domain II, Ala403, Ile407, Gln522, Phe525 and Leu529 were identified as *persistent hubs* across all systems (**Figure 4**). **Figure 4** also shows a selected number of *EC* hub residues in hACE2 sub-domain II that are exclusive to the Omicron sub-lineages at positions 399, 400, 402, 410-412 518, 522, 523, 524, 526, 528, 532, 533, 544, 550, 551, 553 and 554. Previously, it was shown that binding of the RBD to ACE2 results in movement of ACE2 sub-domain II residues towards the catalytic pocket, coinciding with increased carboxypeptidase activity [106]. Here, we identified key allosteric communication residues that could contribute to these structural and functional changes.

In all the Omicron sub-lineages, this communication path was interrupted (**Figure 7**) because of loss of *eigen centrality* at residue positions 326, 355-357, 375, 378 and 379 connecting the hACE2 interface to the zinc binding site. However, compensatory gains in *EC* hubs were noted around the zinc binding site and active site cleft of hACE2 in the Omicron sub-lineages, possibly required to maintain peptidase activity. In BA.1 and BA.3_10, the zinc coordinating residue, Glu402 gained hub status compared to the WT. Remarkably, almost all the residues at positions 326, 355-357, 375, 378 and 379 that lost *EC* hub status experienced higher residue fluctuation in the Omicron sub-lineages compared to the WT (Table S4). This relationship between *EC* and residue flexibility has previously been shown in [75].

Ultimately, the *EC* metric informs the changes in the network patterns of the Omicron sub-lineages resulting from increased residue flexibility leading to a loss of residue influence around the RBM and some RBD neutralizing antibody epitopes. Our findings are in agreement with observations by Cerutti and group, who identified from the Cryo-EM structure of the Omicron S protein, a more structurally dynamic RDB which is believed to elude the recognition and binding by neutralizing antibodies [48]. Increased flexibility in antigenic peptides has been linked to reduced maturation of high affinity antibodies [103].

Furthermore, RBD mutations also affect the inter-protein communication through disruption of the *EC* hub residue network connecting the RBD to the hACE2 active site cleft. The RBD-hACE2 interaction also influences hACE2 carboxypeptidase activity [106] and the loss of network may predict lower influence on hACE2 activity reducing the cardiovascular symptoms related to hACE2 proteolytic activity.

#### 3.5.4 Degree of centrality and Katz centrality

The degree of residue connection in the protein systems was further determined using the *DC* DRN metric and presented as a heat map (**Figure 4**). *DC* ranks residues based on the number of immediate connections. Here, we identified Gly431 and Tyr508 as the only *persistent hubs* in the RBD (**Figure 4 and Figure S7)**. In the WT RBD, *DC* hubs made up the protein core within the α1, α3, β3 and β7 regions of the protein. A general reduction in the RBD *DC* hubs was noted in the majority Omicron sub-lineages compared to the WT probably due to increased dynamics. In the hACE2, *DC* hubs were distributed throughout the structure with residues Ala25, Val93 and Leu97 identified as *DC persistent hubs* (**Figure 4**). Of interest here were residues Gly326, Asp355, Phe356 and Arg357 of hACE2 which showed a significant loss of *DC* exclusively in the Omicron sub-lineages. These residues are positioned at the hACE2 interface with the S RBD where Asp355 and Arg357 form hydrogen bonds and van der Waals contacts with Thr500 of the RBD, respectively. Mapping of the *DC* hubs onto the 3D structure showed minimal differences in hub distribution between the WT and Omicron sub-lineage systems (**Figure S7**).

With *katz centrality* (*KC*), residues Ile402, Tyr508 and Val510 were identified as *persistent hubs* in the RBD and, Ala25, Leu97, Ala403, Ile407, Gln522, Phe525, Leu529 and Ala550 as *persistent hubs* in hACE2 (**Figure 4**). Similar distribution patterns were noted between the *DC* and *KC* hubs since *KC* determines the relative influence of a node taking into account the immediate and non-immediate neighbors [133]. Here too, residues Gly326, Asp355, Phe356 and Arg357 including Leu351 did not have hub status in any of the Omicron sub-lineages. However, compensatory gains in centrality were noted for BA.2 and BA.3_12 hACE2 interface residues Y41 and Q42 involved in inter-protein interactions with the S RBD (**Figure S8**).

The observed loss and compensatory gains in centrality at key hACE2 interface residues highlight that 1) the RBD mutational effect on centrality is not limited to the RBD but crosses over to the hACE2, and 2) the myriad of Omicron RBD mutations inevitably affect the RBD-hACE2 binding potential. The key RBD-hACE2 interactions were further investigated in the next section.

### 3.6 SARS-CoV-2 Omicron sub-lineages experienced more RBD-hACE2 interactions compared to the WT

Inter-protein (RBD-hACE2) interaction differences between the WT and RBD Omicron sub-lineages over the simulation period were investigated using the *contact_map.py* script from MDM-TASK web [88]. RBD-hACE2 interface residues were identified in the WT structure (PDB ID: 6M0J) through alanine scanning using the ROBETTA webserver [93]. Contact map analysis was performed per system for every RBD interface residue, namely Lys417, Ile418, Tyr449, Tyr453, Leu455, Phe456, Ser477, Phe486, Asn487, Tyr489, Gln493, Gln498, Thr500, Asn501, Val503 and Tyr505. Subsequently, sub-lineage residue contact weights were compared to the WT through delta calculations (WT contact weights – sub-lineage contact weights) to determine mutation-imposed changes on inter-protein interactions (**Figure 8**). In **Figure 8A,** blue implies higher contact frequency between a given pair of RBD-hACE2 residues in the sub-lineage system compared to the WT, white means no contact difference and red implies reduced contact frequency in the sub-lineage system compared to the WT.

**Figure 8:**
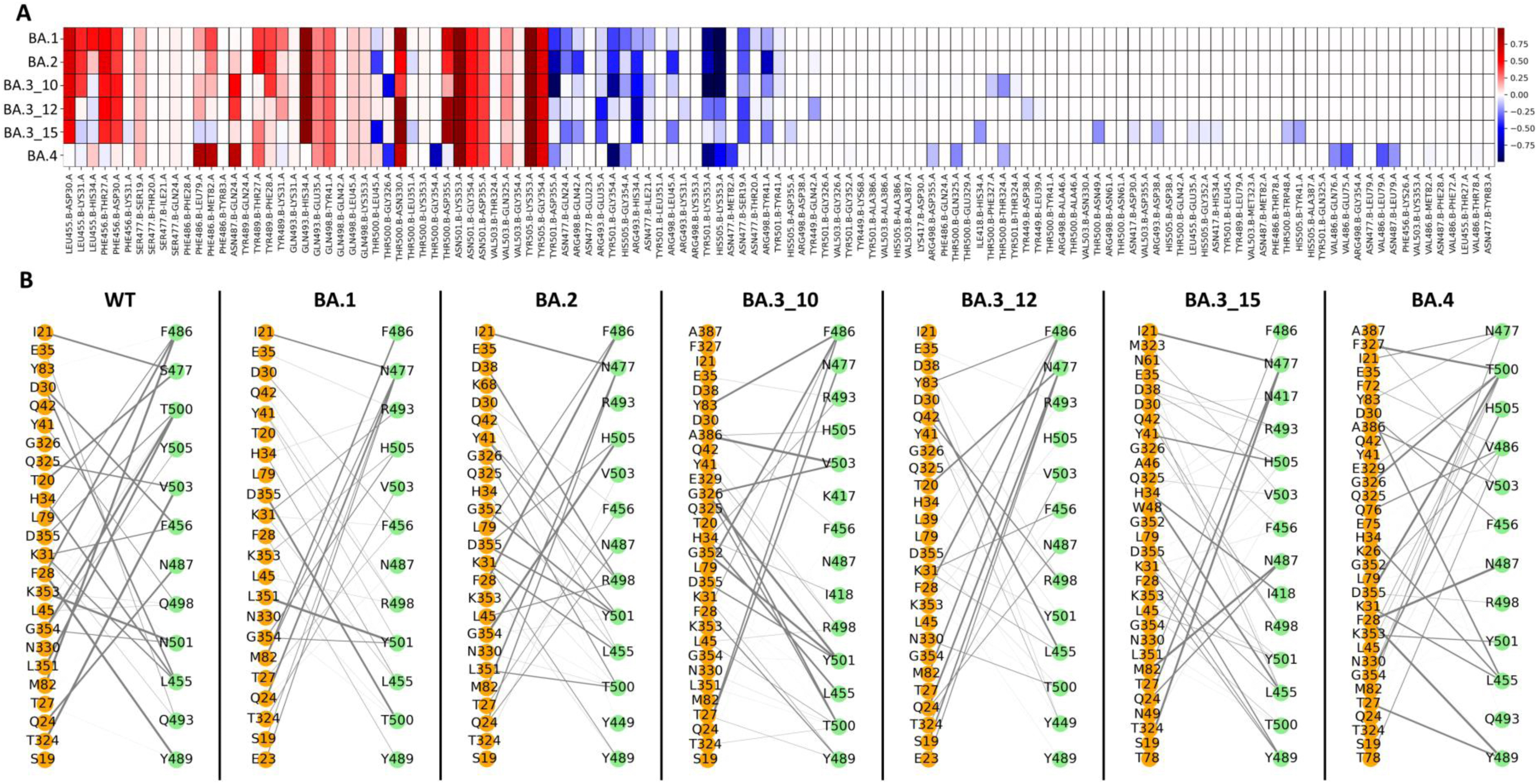
A) Heat map of the delta RBD-hACE2 residue contact frequencies between the WT and the Omicron sub-lineages (WT – sub-lineage). Blue shows higher contact frequency in the sub-lineage to the WT between a given RBD-hACE2 residue pair, white shows no difference and red shows lower residue pair contact frequency in the sub-lineage. B) WT and Omicron sub-lineage RBD-hACE2 interface residue contacts are represented as networks. The orange and green nodes represent the hACE2 and RBD interface residues, respectively, and the weighted grey lines connecting them show the contact frequency.

We noted a general decrease in contact frequency (**Figure 8A**, in red) between the following pairs of RBD-hACE2 protein interface residues in all Omicron sub-lineages compared to WT: Gln493-Glu35, Gln498-Tyr41, Thr500-Asp30, Asn501-Lys353, Asn501-Gly354, Asn501-Asp355, Val503-Gln325, Tyr505-Lys353, and Tyr505-Gly354. Contact frequencies decreased in the following pairs in all Omicron lineages, except BA.4: Leu455-Asp30, Phe456-Thr27, Phe456-Asp30, Gln493-His34, and Thr500-Asp355. We also identified contacts unique to BA.4 sub-lineage, namely Phe486-Leu79, Phe486-Met82 and Asn487-Gln24.

Interestingly, an increase in contact frequency (**Figure 8A**, in blue) between the following pairs of RBD-hACE2 protein interface residues in BA.4 Omicron sub-lineage and in some others was noted compared to WT: Gly447-Met82, Thr500-Thr324, Thr500-Gly326, Asn501-Lys353, Asn501-Asp355, Asn501-Gly354, Tyr505-Gly354, Tyr505-Lys353. The ones unique to BA.4 included Phe486-Glu75, Phe486-Gln76, Phe486-Leu79, Asn487-Leu79, Thr500-Gln325, Thr500-Gly354.

For the RBD mutation at residue 501, which has a documented effect of increased S protein binding affinity [124, 125, 134], more interactions were noted in the Omicron sub-lineages compared to the WT. That is, the WT had 3 hACE2 interactions at that position (Lys353, Gly354 and Asp355), while BA.1 and BA.2 had 5 interactions (Tyr41, Leu351, Lys353, Gly354 and Asp355), BA.3_10 had 6 interactions (Tyr41, Gly326, Gly352, Lys353, Gly354 and Asp355), BA.3_12 had 4 interactions (Tyr41, Lys353, Gly354 and Asp355), BA.3_15 had 4 interactions (Leu45, Gly352, Lys353 and Asp355) and BA.4 had 6 interactions (Tyr41, Gln325, Gly352, Lys353, Gly354 and Asp355). Furthermore, from the network presentation of the complex interactions (**Figure 8B**), it is evident that most of the Omicron sub-lineages accommodate more RBD-hACE2 cross protein interactions compared to the WT i.e., WT: 24, BA.1: 22, BA.2: 25, BA.3_10: 30, BA.3_12: 26, BA.3-15: 30 and BA.4: 32.

Several new RBD-hACE2 interactions were identified in the Omicron sub-lineages during the simulation compared to the WT. These were BA.2: Arg498-Asp38, Tyr449-Lys68 and Tyr501-Gly352; BA.3_10: Thr500-Phe327, Thr500-Glu329, Tyr501-Gly352 and Val503-Ala386; BA.3-12: Asn477-Glu23, Arg498-Asp38, Tyr499-Asp38 and Tyr499-Leu39; BA.3_15: Val503-Met323, Arg498-Asn61, Thr500-Asn61, Arg493-Asp38, His505-Asp38, Arg498-Ala46, Thr500-Ala46, Thr500-Trp48, Tyr501-Gly352, His505-Gly352, Thr500-Asn49 and Phe486-Thr78 and BA.4: Val503-Ala387, His505-Ala387, Val486-Phe72, Val503-Ala386, Thr500-Glu329, Val486-Gln76, Val486-Glu75, Phe456-Lys26, Tyr501-Gly352 and Val486-Thr78. Furthermore, some RBD-hACE2 interactions were more frequent in the Omicron sub-lineages compared to the WT as shown in **Figure 8B**.

It is evident from the interaction analysis that, in addition to the changes in residue centrality especially in *CC* and complex interaction distance, dynamicity of the Omicron sub-lineages predict better binding to the hACE2 host receptor compared to the WT. Prior studies have also shown that Omicron mutations in the RBD facilitate improved binding to the hACE2 compared to the WT virus [135, 136].

## Conclusion

This study aimed to characterize the collective influence of mutations in Omicron sub-lineages, BA.1, BA.2, BA.3 and BA.4, on the RBD-hACE2 interaction, as well as on the behavior of the individual protein domains (RBD and N-terminal of hACE2) using combined computational approaches, including DRN analysis [75–78].

From a global perspective, the RBD of the Omicron sub-lineages sampled a more diverse conformational space than the WT as per ED, RMSD, and Rg calculations. RMSF calculations attributed the diverse conformational nature of the RBD to the highly flexible RBM. The dynamic nature of the RBD also influenced the overall complex dynamics affecting the hACE2. Antigenic hot spots known for binding neutralizing antibodies had greater residue fluctuation in the Omicron sub-lineages which could represent an antibody escape mechanism. Furthermore, the Omicron sub-lineages experienced anti-correlated motions between the RBD and hACE2 proteins, characterized by increased inter-protein COM distance. Protein-focused DCC calculations suggested that the flexible RBM was the cause of the observed anti-correlated motions in the Omicron sub-lineage RBD, especially in BA.1, BA.2 and BA.3_12, and which was also reflected in the hACE2 protein. Previous studies have highlighted the increased binding affinity of the Omicron sub-lineage RBD to the receptor hACE2 protein compared to the reference virus [124, 126, 136, 137]. Here we hypothesize that the observed RBM flexibility favors increased interactions between the S RBD and the receptor hACE2. This was investigated through local residue analysis.

Residue level analyses of the RBD-hACE2 complexes using the MDM-TASK-web [88, 89] highlighted increased RBD-hACE2 residue interactions and interaction frequency of the Omicron sub-lineages compared to the reference protein. Furthermore, centrality metrics of DRN identified for the first time, key allosteric communication pathways between the RDB and hACE2, and evolutionary changes in these networks in the Omicron sub-lineages. *BC* metric provides information on the control of information flow. Strikingly, we identified two allosteric communication paths (Path I and II) in the WT formed by the high centrality *BC* hubs, one of which originated from the RBD core, traversing the receptor binding motif of the S protein and the N-terminal domain of the hACE2, to the active site. We also observed drastic changes in this allosteric path (Path I) while the virus evolved from BA.1 to BA.4. The most dramatic changes were observed in the BA.3 sub-lineages, while in BA.4 the allosteric path was becoming similar to that of the reference protein’s path. This indicates that while the RBD became more flexible in BA.4 via new mutations, the RBD also partially preserved the information flow path in the reference protein.

Increased inter-protein interaction distance was associated with reduced *CC* of the RBD interfacial residues. More so, a depreciating effect of *CC* hubs was noted in the BA.3 sub-lineage sequences as the number of RBD mutations increased.

The *EC* calculations showed a profound reduction in centrality in the Omicron sub-lineages attributed to increased RBD flexibility compared to the WT. Interestingly, this effect translated to the hACE2 protein *EC*. Here, a network of residues previously shown to bridge the RBD to the zinc-binding domain lost *EC* hub status in the Omicron sub-lineages, creating a break in the network chain. Previous work by Lu & Sun showed that binding of the reference S RBD to the hACE2 led to an up to tenfold increase in proteolytic activity of the hACE2 [106]. Based on *BC* and *EC* results, we hypothesize that S RBD mutations affect the peptidase activity of the hACE2 [106].

Taken together, this study presented novel insight, particularly on the evolutionary behavior of the Omicron sub-lineages, where the virus mutated in stages to improve interaction with the receptor, whilst simultaneously retaining critical functional features (e.g. the communication between the viral protein and human receptor). These findings are highly informative for COVID-19 drug and vaccine design.

## Associated Content

### Author Contributions

Conceptualization – ÖTB; Formal analysis – VB and ÖTB; Funding acquisition – ÖTB; Methodology – VB and ÖTB; Project administration – ÖTB; Resources – ÖTB; Supervision –ÖTB; Visualization –VB and ÖTB; Writing - original draft and review & editing – VB, ALE and ÖTB.

## Funding

This work was supported by Funding for COVID-19 Research and Development Goals for Africa Programme (Grant number: SARSCov2-2-20-002) of the African Academy of Sciences (AAS). It is implemented through the Alliance for Accelerating Excellence in Science in Africa (AESA) platform, an initiative of the AAS and the African Union Development Agency (AUDA-NEPAD). It was also supported by the South African Medical Research Council under a Self-Initiated Research Grant awarded to A.L.E. The funders had no role in study design, data collection and analysis, decision to publish, or preparation of the manuscript. The content of this publication is solely the responsibility of the authors and does not necessarily represent the official views of the funders.

### Notes

The authors declare no competing financial interest.

### Data and Software Availability

All data reported in this article are presented in the article and the Supporting Information section. Dynamic residue network analysis metric scripts are implemented in the MDM-TASK-web platform (https://mdmtaskweb.rubi.ru.ac.za/) and are available at https://github.com/RUBi-ZA/MD-TASK/tree/mdm-task-web. MD simulations will be made available upon request.

## Acknowledgement

Authors acknowledge the use of the Centre for High Performance Computing (CHPC), Cape Town, South Africa, for the molecular dynamics simulations.

## Notes

### Competing Interest Statement

The authors have declared no competing interest.

