## Supplementary material for "Evolutionary progression of collective mutations in Omicron sub-lineages towards efficient RBD-hACE2: allosteric communications between and within viral and human proteins": SD

**Table S1:** Omicron sub-lineage specific GISAID sequence IDs

| Omicron sub-lineages | GISAID sequence IDs | |
|---|---|---|
| BA.1 | EPI_ISL_11765237 | EPI_ISL_8097234 |
| | EPI_ISL_11765238 | EPI_ISL_8097272 |
| | EPI_ISL_11765240 | EPI_ISL_8826405 |
| | EPI_ISL_8826449 | EPI_ISL_8826412 |
| | EPI_ISL_9002773 | EPI_ISL_8826429 |
| | EPI_ISL_9002788 | EPI_ISL_7834433 |
| | EPI_ISL_9002795 | EPI_ISL_7834437 |
| | EPI_ISL_9002810 | EPI_ISL_7834440 |
| | EPI_ISL_9002815 | EPI_ISL_7834441 |
| | EPI_ISL_9002836 | EPI_ISL_7834442 |
| | EPI_ISL_7834473 | EPI_ISL_7834443 |
| | EPI_ISL_7834476 | EPI_ISL_7834444 |
| | EPI_ISL_7834514 | EPI_ISL_7834446 |
| | EPI_ISL_7834572 | EPI_ISL_7834453 |
| | EPI_ISL_7834591 | EPI_ISL_7834457 |
| | EPI_ISL_7566360 | EPI_ISL_7552702 |
| | EPI_ISL_7834419 | EPI_ISL_7566330 |
| | EPI_ISL_7834432 | |
| BA.2 | EPI_ISL_10981812 | |
| | EPI_ISL_8636462 | |
| | EPI_ISL_9149786 | |
| | EPI_ISL_10893972 | |
| | EPI_ISL_10967591 | |
| BA.3 | EPI_ISL_7834448 | |
| | EPI_ISL_7852875 | |
| | EPI_ISL_9149817 | |
| | EPI_ISL_9313505 | |
| BA.4 | EPI_ISL_11674413 | |
| | EPI_ISL_11674425 | |
| | EPI_ISL_11674426 | |
| | EPI_ISL_11674430 | |
| | EPI_ISL_11674431 | |
| | EPI_ISL_11674432 | |
| | EPI_ISL_11674443 | |
| | EPI_ISL_11674447 | |
| | EPI_ISL_1167444 | |
| | EPI_ISL_12119232 | |
| | EPI_ISL_11674410 | |
| | EPI_ISL_11674411 | |

**Table S2**. RBD mutations per Omicron sub-lineage

| Sub-lineage | RBD mutations |
|---|---|
| BA.1 | G339D, S371L, S373P, S375F, K417N, N440K, G446S, S477N T478K, E484A, Q493R, G496S, Q498R, N501Y, Y505H |
| BA.2 | G339D, S371F, S373P, S375F, T376A, D405N, R408S, K417N, N440K, S477N, T478K, E484A, Q493R, Q498R, N501Y, Y505H |
| BA.3 | G339D, S373P, S375F, S477N, T478K, E484A, Q493R, Q498R, N501Y, Y505H |
| | G339D, S371L, S373P, S375F, S477N, T478K, E484A, Q493R, G496S, Q498R, N501Y, Y505H |
| | G339D, S371F, S373P, S375F, D405N, K417N, N440K, G446S, S477N, T478K, E484A, Q493R, Q498R, N501Y, Y505H |
| BA.4 | G339D, S371F, S373P, S375F, T376A, D405N, R408S, K417N, N440K, L452R, S477N, T478K, E484A, F486V, Q498R, N501Y, Y505H |

**Table S3**: Residue physicochemical properties

| Mutation | Physicochemical property change |
|---|---|
| G446S | Hydrophobic - Hydrophilic |
| S371F | Small/Polar - Hydrophobic |
| G339D | Non-polar - Polar |
| Q498R | Polar/uncharged – Polar/+ve charge |
| F486V | Hydrophobic - Hydrophobic |
| Y505H | Aromatic - Aromatic |
| T376A | Hydrophobic - Hydrophobic |
| S371L | Small - Aliphatic |
| Q493R | Polar/uncharged - Polar/+ve charge |
| D405N | Small/charged -Small |
| N501Y | Small – Aromatic |
| R408S | +ve charge - Polar |
| S477N | Polar/tiny - Polar |
| G496S | Small - Small/Polar |
| S373P | Small/Polar - Small |
| T478K | Polar – Polar/+ve charge |
| E484A | Polar - Nonpolar |
| S375F | Polar/small - Hydrophobic/Aromatic |
| L452R | Hydrophobic/aliphatic - Polar /+ve charge |
| K417N | Polar - Polar |
| N440K | Polar - Polar |

**Table S4**: *EC* hub residue flexibility

| | RMSF (nm) | | | | | | |
|---|---|---|---|---|---|---|---|
| **Residue** | **WT** | **BA.1** | **BA.2** | **BA.3_10** | **BA.3_12** | **BA.3_15** | **BA.4** |
| Gly326 | 0.192 | 0.215 | 0.218 | 0.141 | 0.255 | 0.206 | 0.184 |
| Asp355 | 0.096 | 0.113 | 0.169 | 0.097 | 0.221 | 0.153 | 0.127 |
| Phe356 | 0.096 | 0.121 | 0.176 | 0.099 | 0.227 | 0.164 | 0.134 |
| Arg357 | 0.103 | 0.136 | 0.196 | 0.117 | 0.236 | 0.179 | 0.147 |
| Glu375 | 0.107 | 0.147 | 0.194 | 0.140 | 0.254 | 0.162 | 0.145 |
| His378 | 0.092 | 0.120 | 0.154 | 0.111 | 0.154 | 0.153 | 0.123 |
| Ile379 | 0.085 | 0.112 | 0.145 | 0.123 | 0.186 | 0.192 | 0.122 |

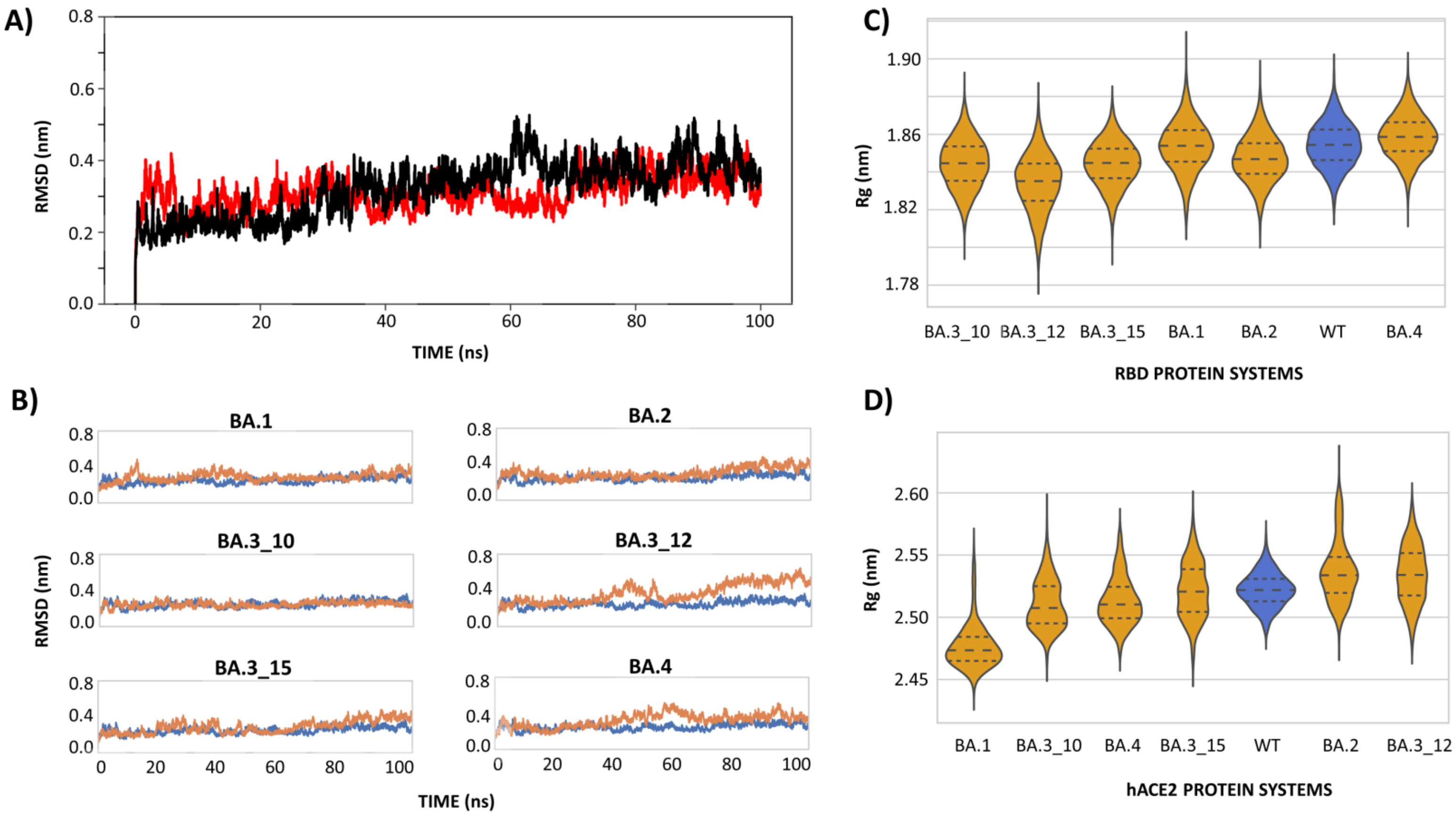


**Figure S1**: A) RBD-hACE2 WT quality control RMSD distribution runs one (red) and two (black). B) Progressive RMSD line plots comparing the conformational evolution of the WT (blue) to the Omicron sub-lineages (orange) for each RBD-hACE2 protein complex. C and D show the Rg violin plot distribution of the WT (blue) and mutant systems (orange) in the RBD and hACE2 proteins, respectively. The x-axis displays the protein systems whereas the y-axis shows the Rg values in both sub plots.

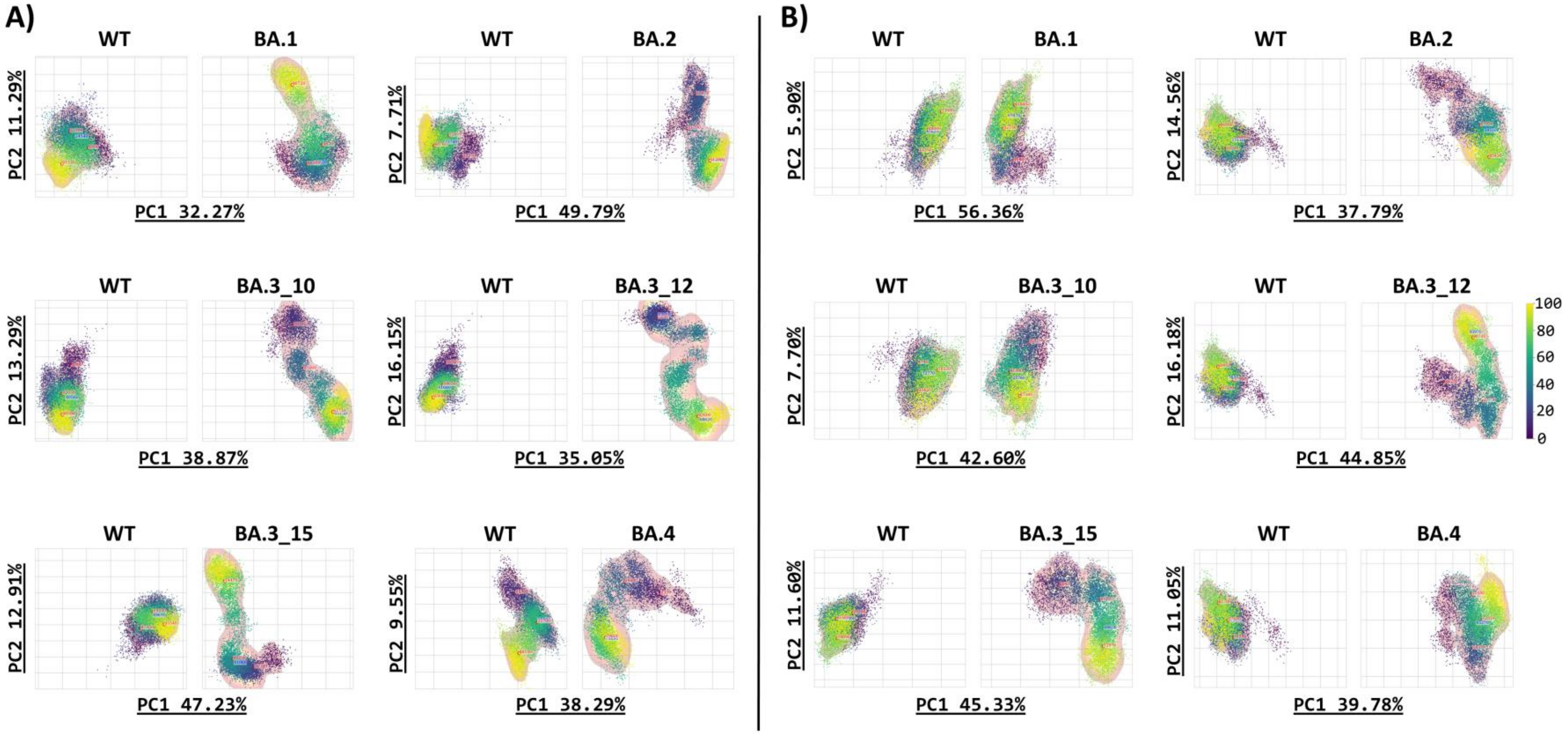


**Figure S2**: Scatter plots of comparative essential dynamics of the A) RBD and B) hACE2 in the WT and Omicron sub-lineage systems. The WT is left in each subplot and the Omicron sub-lineage system to the right. The x- and y-axes show the variance in percentage as explained by PC1 and PC2, respectively.

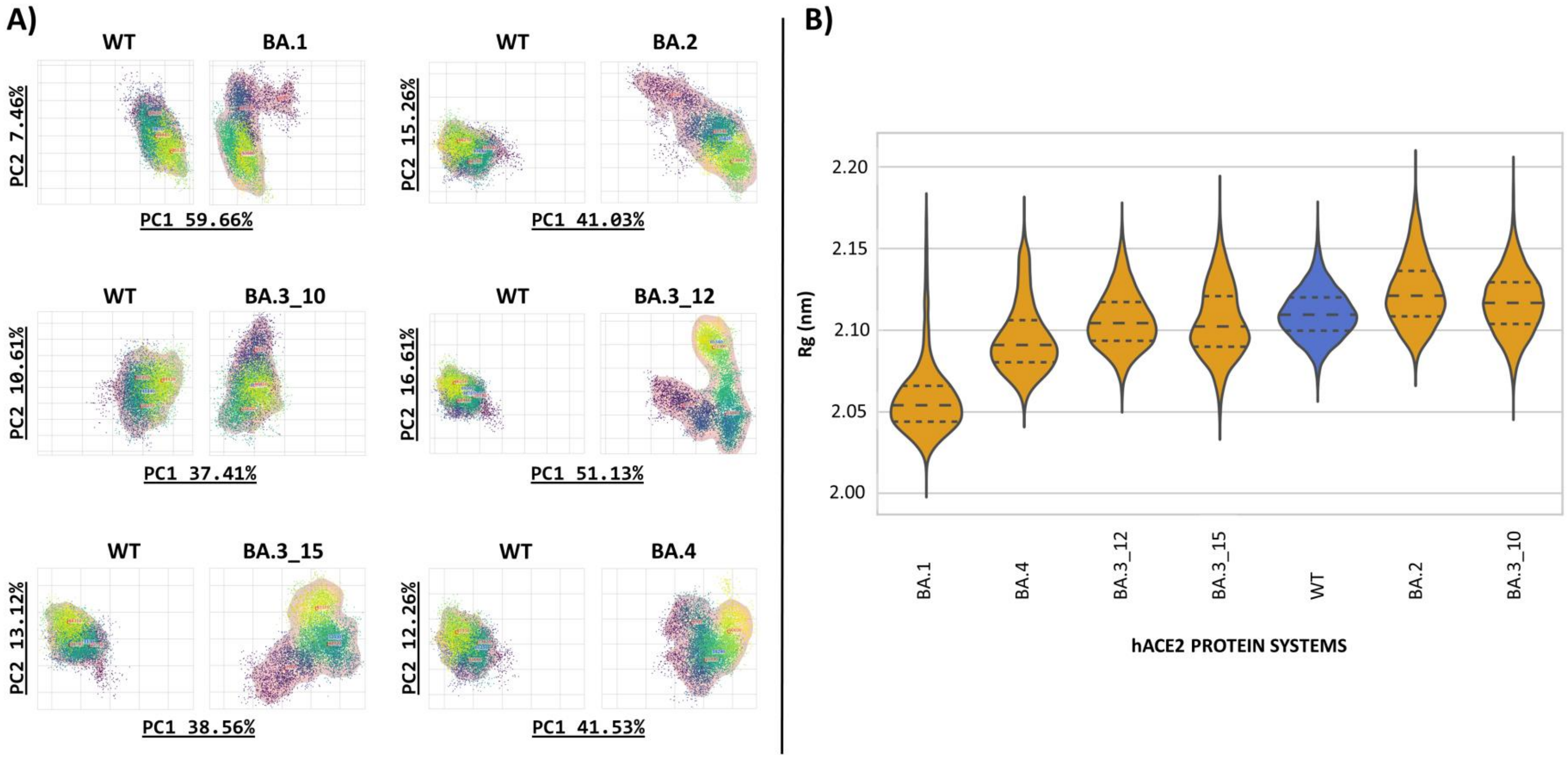


**Figure S3**: A) Scatter plots of comparative essential dynamics of the hACE2 groove cavity in the WT and Omicron sub-lineage systems. The WT is left in each subplot and the Omicron sub-lineage system to the right. The x- and y-axes show the variance in percentage as explained by PC1 and PC2, respectively. B) Violin plots of the WT (blue) and Omicron sub-lineage (orange) hACE2 substrate binding cleft Rg. The x-axis displays the protein systems whereas the y-axis shows the Rg values.

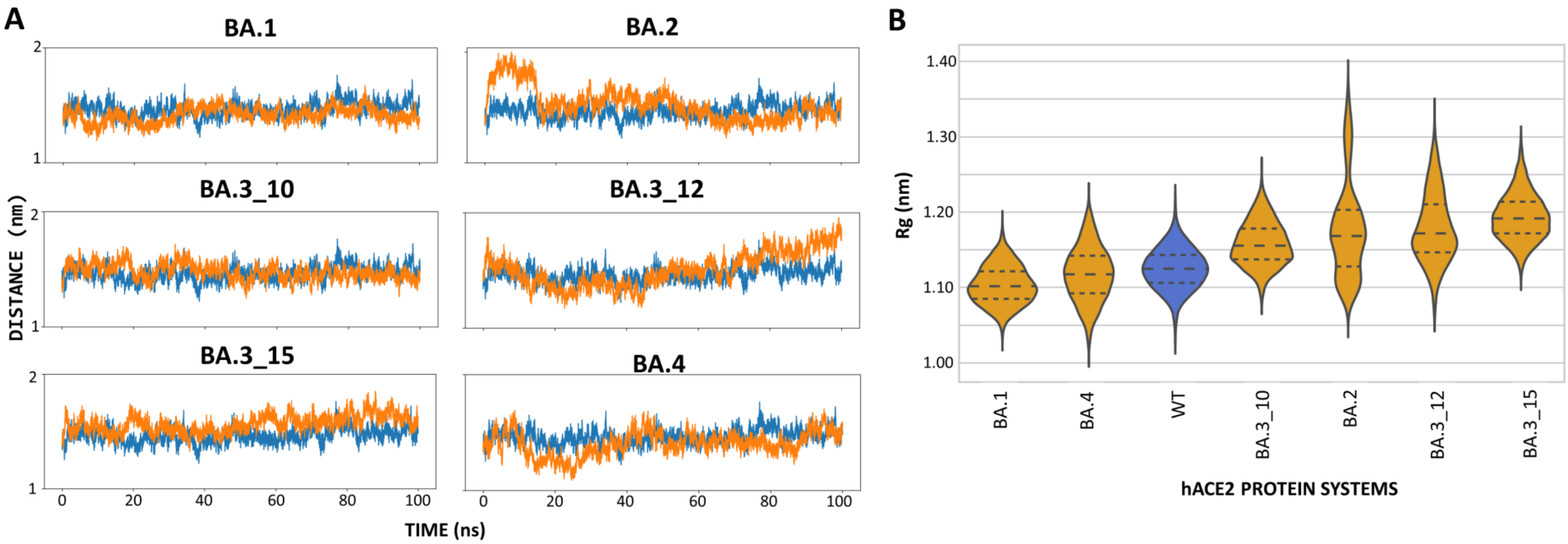


**Figure S4**: A) Comparative residue COM distance between the hACE2 active site residues in sub-domain I and II for the WT (blue) and each Omicron sub-lineage (orange). The x- and y-axes show the time (ns) and distance (nm), respectively. B) Violin plots of the WT (blue) and Omicron sub-lineage (orange) hACE2 active site residue Rg arranged in ascending order of the median Rg. The x-axis displays the protein systems whereas the y-axis shows the Rg values.

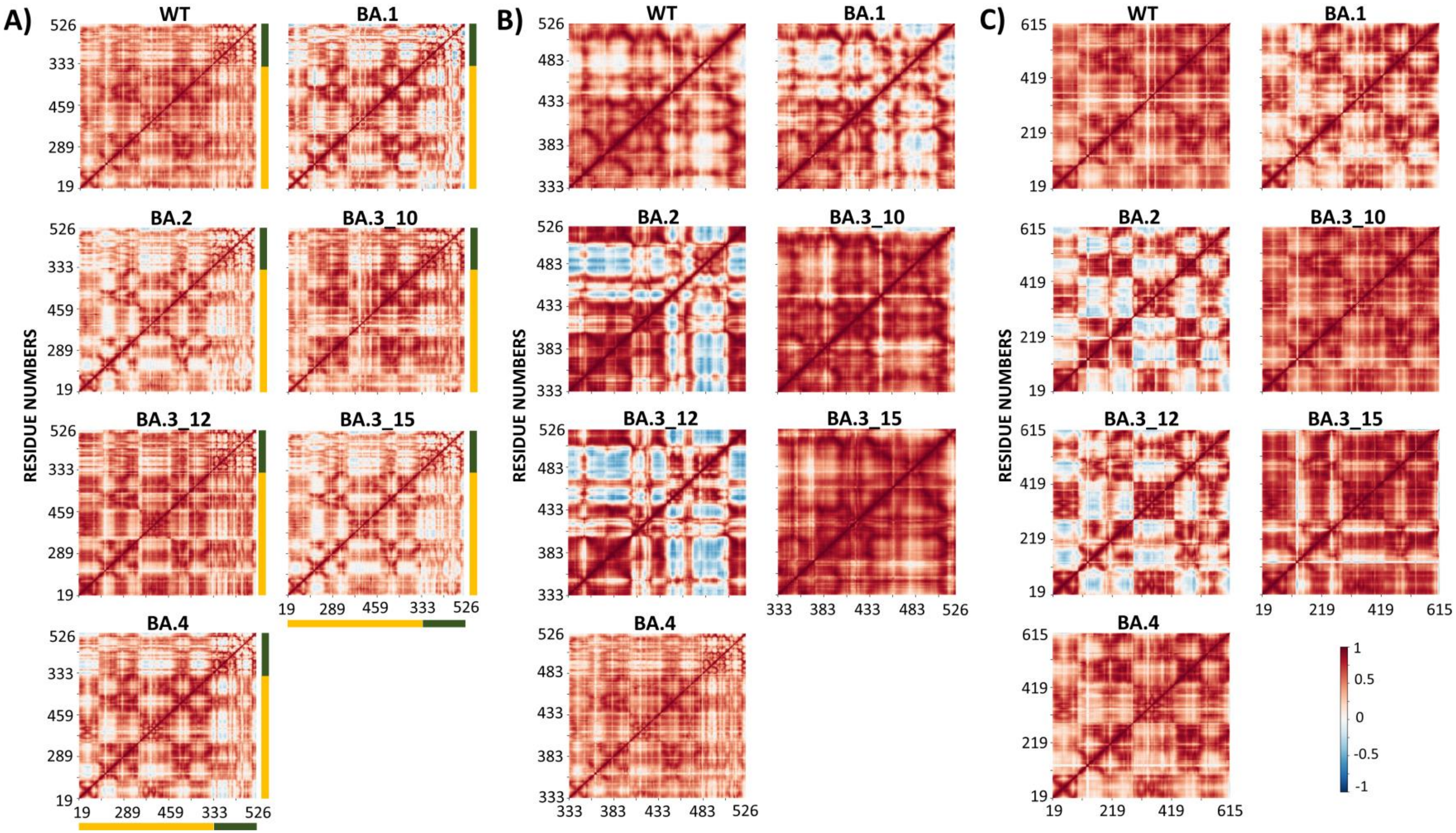


**Figure S5**: Heat maps of the degree of DCC in the A) RBD-hACE2 complexes, B) RBD and C) hACE2 for the WT and Omicron sub-lineages. The scale from 1 through 0 to -1 indicates the degree of correlation, with 1 being highly correlated, 0 no correlation, and -1 highly anti-correlated motions. The orange and green heatmap sections mark the beginning and end of the hACE2 and RBD protein sections, respectively.

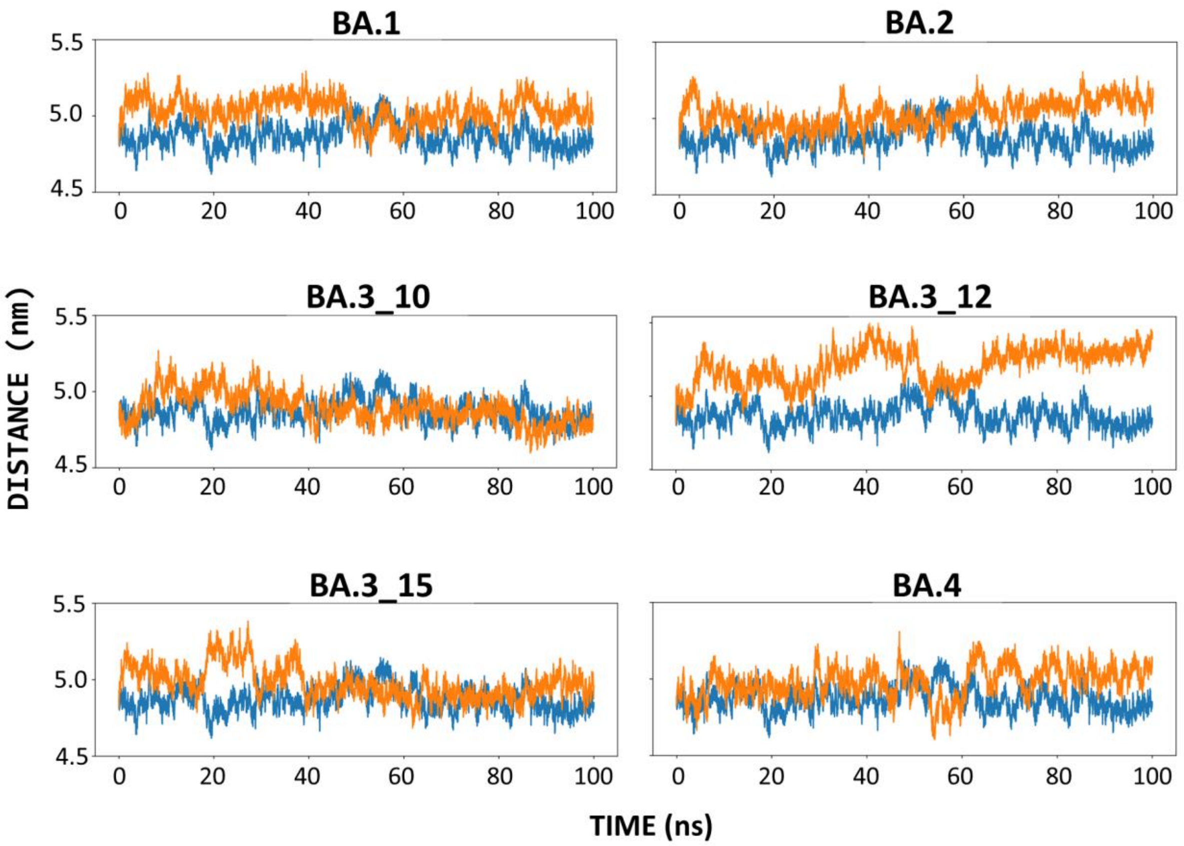


**Figure S6**: Comparative RDB-hACE2 COM distance between the WT (blue) and each Omicron sub-lineage (orange). The x- and y-axes show the time (ns) and distance (nm), respectively.

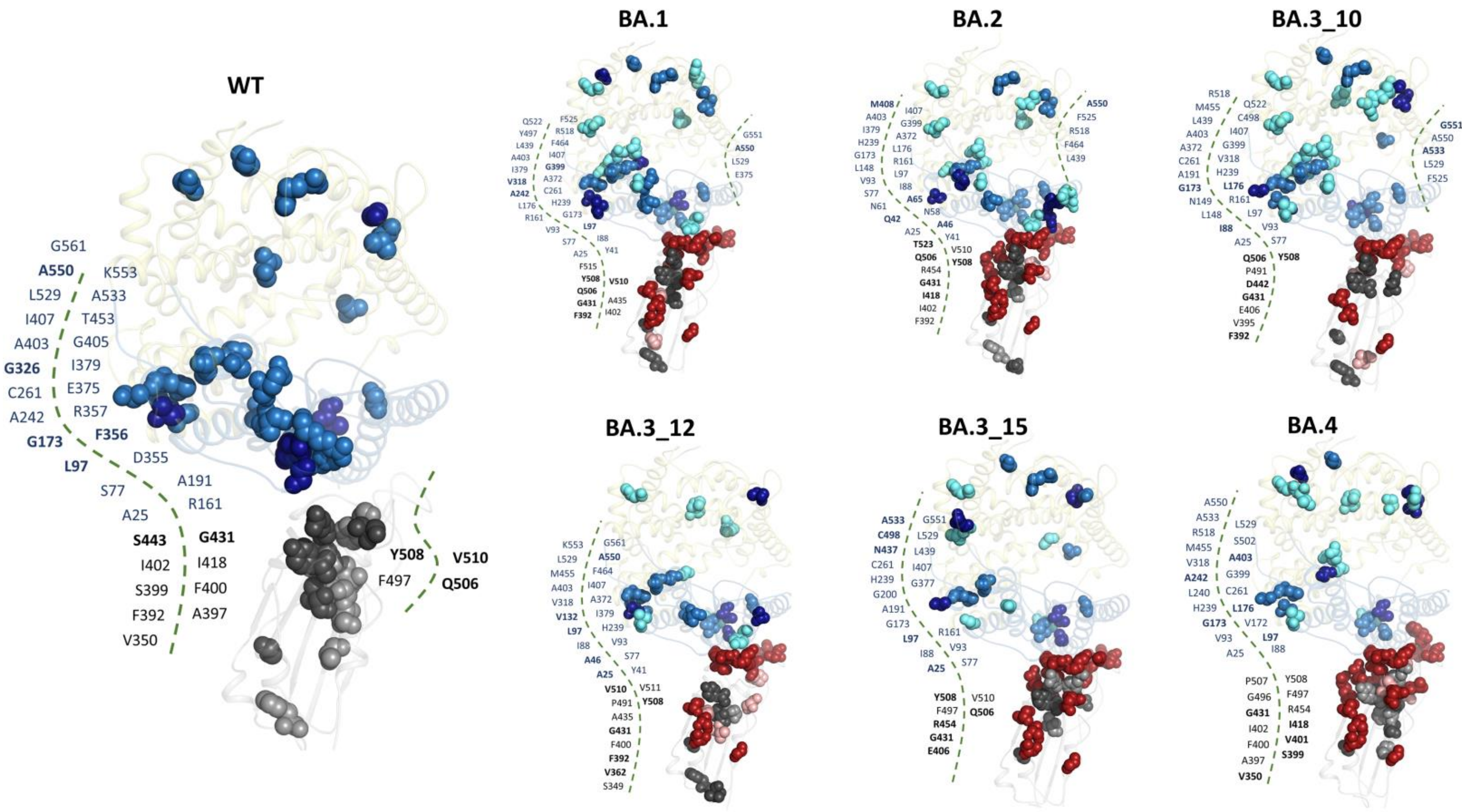


**Figure S7**: Cartoon representation of the RBD-hACE2 structures showing the distribution of the global top 5% and 4% *DC* hubs in the RBD and the hACE2, respectively for the WT and omicron sub lineages. WT hubs are shown as sky-blue spheres (hACE2) and grey spheres (RBD). The same colors are used for *DC* hubs common to the WT and omicron variants. *DC* hubs unique to the variants (Δ hubs: mutant hubs – WT hubs) are shown as aquamarine spheres (hACE2) and boron spheres (RBD). The five highest centrality residues in RBD and hACE2 are shown as dark grey and dark blue spheres, respectively, and annotated in bold. The variant specific mutation positions are shown as firebrick spheres.

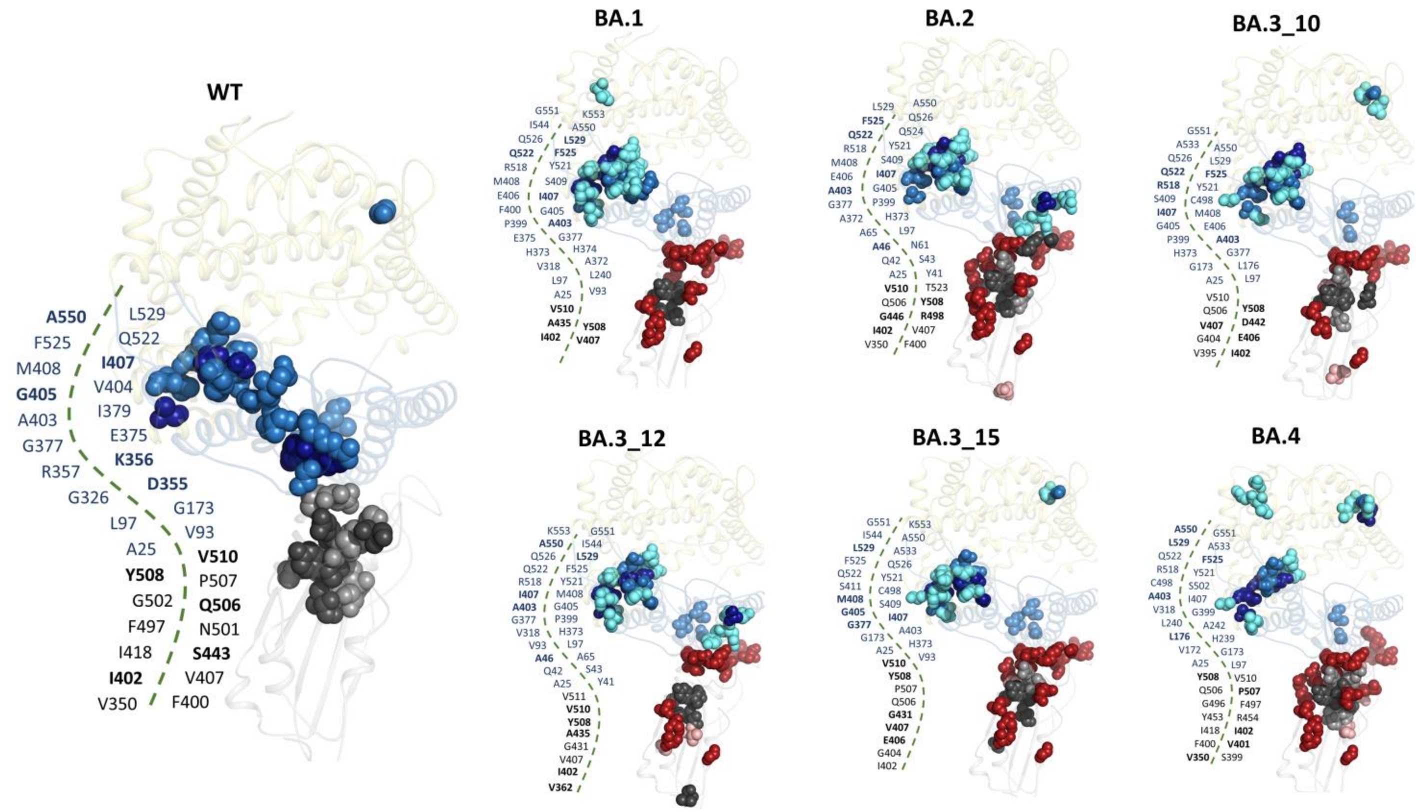


**Figure S8**: Cartoon representation of the RBD-hACE2 structures showing the distribution of the global top 5% and 4% *KC* hubs in the RBD and the hACE2, respectively for the WT and omicron sub lineages. WT hubs are shown as sky-blue spheres (hACE2) and grey spheres (RBD). The same colors are used for *KC* hubs common to the WT and omicron variants. *KC* hubs unique to the variants (Δ hubs: mutant hubs – WT hubs) are shown as aquamarine spheres (hACE2) and boron spheres (RBD). Interface hACE2 hub residues exclusive to the WT are labeled in black and the compensatory gains in centrality in the variants are labeled in blue. The five highest centrality residues in RBD and hACE2 are shown as dark grey and dark blue spheres, respectively, and annotated in bold. The variant specific mutation positions are shown as firebrick spheres.
